## Supplementary Information for "Directing Min protein patterns with advective bulk flow"


##### Code and data availability

The simulation and experimental data generated for this study can be found on a public repository on Zenodo at https://zenodo.org/record/6818110.

#### 1 Theoretical Methods

In the following, we summarize the theoretical methods used in this paper. The structure is as follows: In Sec. 1.1 we present the theoretical model of the experimental setup – this amounts to extending the *switch model* introduced in Ref.[4] to account for advective bulk flow. As numerical simulations of the full system are costly, we justify and implement several simplifications of the full geometry (Sec. 1.2). Next, we show in Sec. 1.3 how in the limit of high MinE:MinD ratio, the reaction kinetics of the switch model can be simplified, arriving at the *reduced switch model*. We then state the equations of motion and parameters for the switch model (Sec. 1.4.1), the reduced switch model (Sec. 1.4.2), and the *skeleton model* (Sec. 1.4.3), which does not include conformational switching of MinE. We then describe how the effects of advective flow on the presented models are analysed by means of finite element simulations and a linear stability analysis. Finally, in Sec. 1.8 we present three other theoretical models from the literature[1,5,8], and extend them to include molecular switching of MinE. We conclude by summarizing the findings of a preliminary analysis of the effects of flow on these models.

##### 1.1 Setup

In order to simulate the full experimental system, one would have to perform Finite Element Method (FEM) simulations in a 2+3D box geometry, with 2-dimensional membranes at the top and bottom encapsulating a 3-dimensional box of cytosol (see panel A in Fig. S1). Since we are predominantly interested in phenomena that occur in the direction of flow, we consider as a starting point a 1+2D slice of the full 2+3D geometry, where the direction of "slicing" is parallel to the direction of advective flow.

The slice geometry (panel B in Fig. S1) is a rectangle $[0, L] \times [0, H]$, where $L$ is the length of the slice and $H$ is the height of the bulk. The membrane is located at $z = 0$ In this geometry, the switch model can be formulated as follows, starting with the membrane species:

$$\partial_t m_{\mathrm{d}}(x,t) = D_m \partial_x^2 m_{\mathrm{d}} + f_d \tag{1a}$$

$$\partial_t m_{\mathrm{de}}(x,t) = D_m \partial_x^2 m_{\mathrm{de}} + f_{\mathrm{de}}, \tag{1b}$$

where $m_{\mathrm{d}}$ and $m_{\mathrm{de}}$ describe the concentrations of membrane bound MinD and MinDE complexes, respectively. The dynamics of the cytosolic components read:

$$\partial_t c_{\mathrm{DD}}(x,z,t) + \mathbf{v}_f \cdot \nabla c_{\mathrm{DD}} = D_c \nabla^2 c_{\mathrm{DD}} - \lambda c_{\mathrm{DD}} \tag{2a}$$

$$\partial_t c_{\mathrm{DT}}(x,z,t) + \mathbf{v}_f \cdot \nabla c_{\mathrm{DT}} = D_c \nabla^2 c_{\mathrm{DT}} + \lambda c_{\mathrm{DD}} \tag{2b}$$

$$\partial_t c_{\mathrm{Er}}(x,z,t) + \mathbf{v}_f \cdot \nabla c_{\mathrm{Er}} = D_c \nabla^2 c_{\mathrm{Er}} - \mu c_{\mathrm{Er}} \tag{2c}$$

$$\partial_t c_{\mathrm{Ei}}(x,z,t) + \mathbf{v}_f \cdot \nabla c_{\mathrm{Ei}} = D_c \nabla^2 c_{\mathrm{Ei}} + \mu c_{\mathrm{Er}} \tag{2d}$$

Here, $c_{\mathrm{DD}}$ and $c_{\mathrm{DT}}$ denote the cytosolic concentrations of MinD-ADP and MinD-ATP, respectively. $c_{\mathrm{Er}}$ and $c_{\mathrm{Ei}}$ correspond to the concentrations of reactive and latent MinE. Moreover, we used a parabolic flow profile

$$\mathbf{v}_f = \left[v_{\mathrm{f}}^{\max}(h-z)(h+z)/h^2, 0\right],$$

where $v_{\mathrm{f}}^{\max}$ is the flow velocity in the middle of the channel (at $z = 0$).

We are interested in the dynamics far (compared to the wavelength) from the inlet. Therefore, we employ periodic boundary conditions both for the membrane species ($u_m \in \{m_{\mathrm{d}}, m_{\mathrm{de}}\}$)

$$u_m(0,t) = u_m(L,t) \tag{3a}$$

$$\partial_x u_m(0,t) = \partial_x u_m(L,t), \tag{3b}$$

and likewise for the cytosolic species ($u_c \in \{c_{\mathrm{DD}}, c_{\mathrm{DT}}, c_{\mathrm{Er}}, c_{\mathrm{Ei}}\}$):

$$u_c(0,z,t) = u_c(L,z,t) \tag{4a}$$

$$\partial u_c(0,z,t) = \partial_x u_c(L,z,t). \tag{4b}$$

These boundary conditions allow us to study periodic phenomena such as traveling waves on a relatively small domain.

The attachment to and detachment from the membrane are included in the reactive boundary conditions, which couple the membrane and the cytosol:

$$-D_c \partial_z c_{\mathrm{DD}}|_{z=0} = f_{\mathrm{DD}}|_{z=0} \tag{5a}$$

$$-D_c \partial_z c_{\mathrm{DT}}|_{z=0} = f_{\mathrm{DT}}|_{z=0} \tag{5b}$$

$$-D_c \partial_z c_{\mathrm{Er}}|_{z=0} = f_{\mathrm{Er}}|_{z=0} \tag{5c}$$

$$-D_c \partial_z c_{\mathrm{Ei}}|_{z=0} = f_{\mathrm{Ei}}|_{z=0}. \tag{5d}$$

Whereas at $z = H$, we impose a no-flux boundary condition $\partial_z c_i|_{z=H} = 0$ where $i = \mathrm{DD}, \mathrm{DT}, \mathrm{Er}, \mathrm{Ei}$.

The reaction terms, corresponding to the schematic in main text Fig. 1B are:

$$f_{\mathrm{d}} = (k_{\mathrm{D}} + k_{\mathrm{dD}} m_{\mathrm{d}}) c_{\mathrm{DT}} - (k_{\mathrm{dEr}} c_{\mathrm{Er}} + k_{\mathrm{dEi}} c_{\mathrm{Ei}}) m_{\mathrm{d}} \tag{6a}$$

$$f_{\mathrm{de}} = (k_{\mathrm{dEr}} c_{\mathrm{Er}} + k_{\mathrm{dEi}} c_{\mathrm{Ei}}) m_{\mathrm{d}} - k_{\mathrm{de}} m_{\mathrm{de}} \tag{6b}$$

$$f_{\mathrm{DT}} = -(k_{\mathrm{D}} + k_{\mathrm{dD}} m_{\mathrm{d}}) c_{\mathrm{DT}} \tag{6c}$$

$$f_{\mathrm{DD}} = k_{\mathrm{de}} m_{\mathrm{de}} \tag{6d}$$

$$f_{\mathrm{Er}} = k_{\mathrm{de}} m_{\mathrm{de}} - k_{\mathrm{dEr}} m_{\mathrm{d}} c_{\mathrm{Er}} \tag{6e}$$

$$f_{\mathrm{Ei}} = -k_{\mathrm{dEi}} m_{\mathrm{d}} c_{\mathrm{Ei}}, \tag{6f}$$

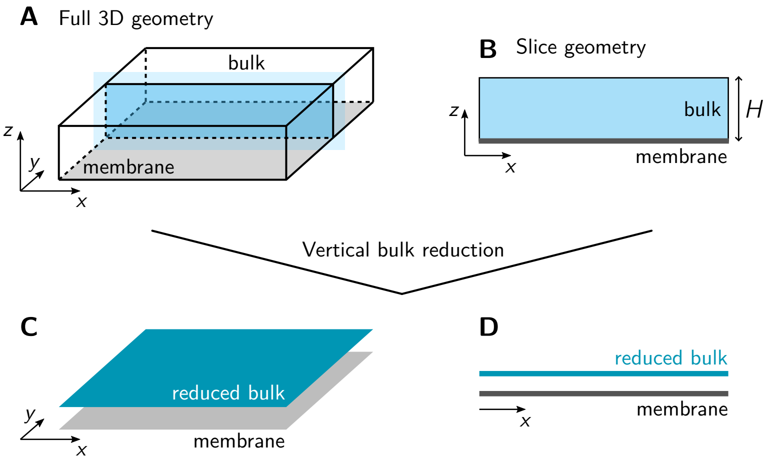


Fig. S1: **A** Bottom half of the full 3D geometry. Blue shaded area corresponds marks the slice visualized in panel B. **B** A 1+2D slice of the full 2+3D geometry. **C** The 2D geometry obtained from the full 3D geometry by eliminating the vertical bulk dimension. **D** The line geometry obtained from the slice geometry by eliminating the vertical bulk dimension.

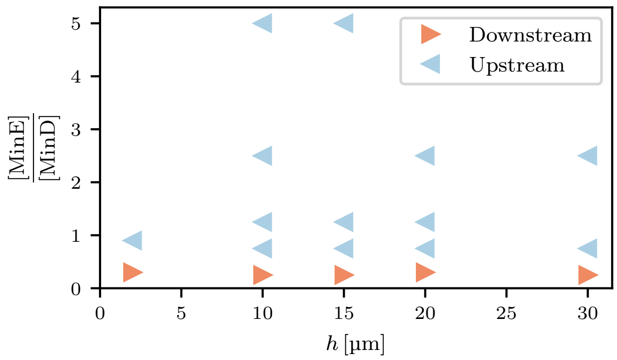


Fig. S2: Results of a parameter sweep of the switch model, where the bulk height $h$ was varied along with the E:D ratio (other parameters can be found in Table S1). Red/blue markers indicate simulations where downstream/upstream propagation was observed.

where the cytosolic components are to be evaluated at the membrane, i.e. at $z = 0$.

Note that the above dynamics conserve the total numbers of MinD and MinE

$$\bar{n}_\mathrm{D} V_\mathrm{bulk} = \int_\mathrm{mem} \mathrm{d}x\, (m_\mathrm{d} + m_\mathrm{de}) + \int_\mathrm{bulk} \mathrm{d}x^2\, (c_\mathrm{DD} + c_\mathrm{DT}), \tag{7}$$

$$\bar{n}_\mathrm{E} V_\mathrm{bulk} = \int_\mathrm{mem} \mathrm{d}x\, m_\mathrm{de} + \int_\mathrm{bulk} \mathrm{d}x^2\, (c_\mathrm{Ei} + c_\mathrm{Er}). \tag{8}$$

Therefore, the average total densities $\bar{n}_\mathrm{D}$ and $\bar{n}_\mathrm{E}$ are important control parameters of the system.

##### 1.2 Eliminating the vertical bulk dimension

Performing simulations in this 2-dimensional slice geometry reveals that bulk height does not significantly affect the wave propagation direction in response to flow (see Fig. S2). Specifically, we find that the E:D ratio where the propagation direction changes from downstream to upstream propagation does not sensitively depend on the the bulk height. This suggests that vertical bulk gradients are not important for the effect of (lateral) advective flow on the wave patterns. We therefore eliminate the vertical bulk dimension by "integrating out" the vertical concentration profiles. This reduces the model to a one-dimensional geometry (see panel D in

Fig. S1), which significantly speeds up numerical simulations. A more detailed discussion of this model reduction is given at the end of this section.

To eliminate the vertical bulk dimension, and thus reduce the model to a one-dimensional (line) geometry, we vertically average the cytosolic concentrations

$$\bar{u}_c(x,t) := H^{-1} \int_0^H \mathrm{d}z\, u_c(x,z,t), \tag{9}$$

for each $u_c \in \{c_{\mathrm{DD}}, c_{\mathrm{DT}}, c_{\mathrm{Ei}}, c_{\mathrm{Er}}\}$. Recall the bulk dynamics

$$\partial_t u_c + v_{\mathrm{f}}(z)\nabla_x u_c = D_c \nabla^2 u_c + \gamma_{u_c}(u_c), \tag{10}$$

where $\gamma_{u_c}(u_c)$ are linear functions of the cytosolic concentrations $u_c$, cf. Eq. (2). We can integrate this equation over the vertical dimension, and divide it by $H$, obtaining

$$\partial_t \bar{u}_c + \bar{v}_f \nabla_x \bar{u}_c = D_c \left[\nabla_x^2 \bar{u}_c + H^{-1}\, \nabla_z u_c|_0^H\right] + \gamma_{u_c}(\bar{u}_c). \tag{11}$$

Here, $\bar{v}_f$ is the average flow velocity in the cytosol. Using the reactive boundary conditions at $z = 0$ (cf. Eq. 5) and the zero-flux boundary condition at $z = H$ yields

$$\partial_t \bar{u}_c + \bar{v}_f \partial_x \bar{u}_c = D_c \partial_x^2 \bar{u}_c + H^{-1} f_{u_c}(u_m, u_c|_{z=0}) + \gamma_{u_c}(\bar{u}_c). \tag{12}$$

To obtain a closed set of equations, we approximate the cytosolic concentrations at the membranes with the vertically averaged concentration as $u_c|_{z=0} \approx \bar{u}_c$ in both the membrane and cytosolic dynamics:

$$\partial_t u_m + \bar{v}_f \partial_x \bar{u}_c = D_m \partial_x^2 u_m + f_{u_m}(u_m, \bar{u}_c) \tag{13a}$$

$$\partial_t \bar{u}_c = D_c \partial_x^2 \bar{u}_c + H^{-1} f_{u_c}(u_m, \bar{u}_c) + \gamma_{u_c}(\bar{u}_c). \tag{13b}$$

By this approximation, we are neglecting vertical concentration gradients in the bulk. This becomes precise if the bulk height is small enough that the bulk diffusion mixes it well on the timescale of the reactions at the membrane. This is not the case in the Min system, unless the bulk height is smaller than about 5 µm (see Brauns, et al.[2]). Indeed, vertical gradients have been shown to play an important role for several characteristics of the Min-protein dynamics in vitro[2,6]. However, as we discussed above, simulations in a domain with vertically extended bulk show no significant effect of the bulk height on the wave propagation direction. Therefore, this *qualitative* effect of bulk flow can be studied in a one-dimensional domain where the bulk is integrated out, as described above. Thus, we study the switch model in a line geometry, with the dynamical equations as in Sec. 1.4.1. This significantly speeds up the simulations and allows us to map out the phase diagram in the parameter plane of flow velocity and E:D ratio. It also allows us to run long simulations and thus perform adiabatic parameter sweeps to demonstrate hysteresis.

##### 1.3 Reducing the switch model

For simplicity, the model reduction is performed for a one-dimensional domain. At the end of this section we briefly discuss how the calculation can be generalized to take into account the extended bulk dimension.

Starting from the one-dimensional dynamics for the switch model as given in Sec. 1.4.1, we consider the limit of high total MinE density ($n_{\mathrm{E}} \to \infty$) and slow recruitment of latent MinE to the membrane ($k_{\mathrm{dEi}} \to 0$), such that their product $k_{\mathrm{dEi}}^{\mathrm{eff}} = k_{\mathrm{dEi}} \cdot n_{\mathrm{E}}$ is kept constant. If conformational switching between reactive and latent MinE is fast ($\mu \to \infty$), the majority of cytosolic MinE is in its latent conformation, forming an effective reservoir. This means we can eliminate the latent cytosolic conformation of MinE $c_{\mathrm{Ei}}$ and replace the terms describing its membrane binding as $k_{\mathrm{dEi}} m_{\mathrm{d}} c_{\mathrm{Ei}} \to k_{\mathrm{dEi}}^{\mathrm{eff}} m_{\mathrm{d}}$.

If we additionally assume that reactive MinE is rapidly recruited to the membrane, we can perform a Quasi Steady State Approximation (QSSA) for the reactive MinE component. Since both processes involving reactive MinE are fast compared to other processes in the system, all potential deviations from its steady state will quickly be neutralized, allowing us to set its reaction term Eq. (18f) to zero. We can thus solve for the QSSA approximation of the reactive MinE concentration:

$$c_{\mathrm{Er}}^{*} \approx \frac{k_{de} m_{\mathrm{de}}}{k_{\mathrm{dEr}} m_{\mathrm{d}} + \mu} \tag{14}$$

Using the obtained results, we can analyze the production of MinDE complexes on the membrane. The effective transition rate between membrane bound MinD and MinDE complexes then reads:

$$k'_{\mathrm{d}\to\mathrm{de}} \approx \left[ k_{\mathrm{dEi}}^{\mathrm{eff}} + \frac{k_{\mathrm{de}} m_{\mathrm{de}}}{\mu / k_{\mathrm{dEr}} + m_{\mathrm{d}}} \right] m_{\mathrm{d}}. \tag{15}$$

The numerator in the second term implies that the formation of MinDE complexes is essentially autocatalytic, since the production term is proportional to $m_{\mathrm{de}}$ itself. The intuitive explanation lies in the fact that MinDE complexes detaching from the membrane are the only source of reactive MinE in the cytosol, which is then quickly recruited back to the membrane by MinD. Omitting this intermediate recruitment step in the limit of rapid membrane recruitment ($k_{\mathrm{dEr}} \to \infty$) then yields the autocatalytic behaviour in Eq. (16).

In the above derivation, we implicitly assumed that the cytosolic concentrations are well-mixed in the direction orthogonal to the membrane. However, this assumption is in conflict with taking the limit of fast MinE deactivation, which results in a sharp gradient of reactive MinE in the vicinity of the membrane.[1] We can account for this by averaging the bulk flux of reactive MinE at the membrane not over the entire domain height $h$ but only over the penetration depth $l_p = \sqrt{D_c/\mu}$. Thus, when averaging the bulk dynamics of reactive MinE, the factor $l_p^{-1}$ replaces the factor $h^{-1}$, since reactions predominantly occur in a thin layer of order $l_p$ in the vicinity of the membrane. Taking this into account yields the corrected transition rate between membrane MinD and MinDE complexes as:

$$k_{\mathrm{d}\to\mathrm{de}} \approx \left[ k_{\mathrm{dEi}}^{\mathrm{eff}} + \frac{k_{\mathrm{de}} m_{\mathrm{de}}}{\sqrt{D_c \mu} / k_{\mathrm{dEr}} + m_{\mathrm{d}}} \right] m_{\mathrm{d}}. \tag{16}$$

This corrected QSSA is valid if we hold the ratio $\sqrt{\mu D_c}/k_{\mathrm{dEr}}$ constant when taking the limits $\mu \to \infty$ and $k_{\mathrm{dEr}} \to \infty$. Indeed, a model reduction in the 2+1D domain where vertical bulk gradients are accounted for yields the same expression for the effective rate $k_{\mathrm{d}\to\mathrm{de}}$.

In essence, we completely eliminated both cytosolic MinE components from the dynamics, arriving at a simplified model of the Min system which we refer to as the *reduced switch model*. The full set of equations that constitute the reduced model is stated in Sec. 1.4.2.

##### 1.4 Summary of models and parameters used

As a consequence of the arguments outlined in Sec. 1.2 and the gained computational simplicity, the simulations presented in this work were performed on a line geometry (see Fig. S1, panel D), namely on an interval of length $L = 500\,\mu\mathrm{m}$, with periodic boundary conditions. Across all simulations, the parameters stated in Tab. S1 were shared. In the following we will use $u(x,t) \equiv u$, i.e. omit explicitly stating the dependencies of the concentrations on the position and time.

[1] In a (2+1) dimensional geometry as presented in Sec. 1.1 the bulk concentration profile of reactive MinE in a laterally uniform steady state is given by $c_{\mathrm{Er}}(z) \propto \cosh\left(\sqrt{\mu/D_c}\, z\right)$.

| Parameter | Value | Description |
|---|---|---|
| $D_c$ | $60\,\mu m^2\,s^{-1}$ | Cytosolic diffusion constant |
| $D_m$ | $0.013\,\mu m^2\,s^{-1}$ | Membrane diffusion constant |
| $\lambda$ | $6\,s^{-1}$ | MinD-ADP to MinD-ATP transition rate |
| $\mu$ | $100\,s^{-1}$ | MinE switching rate |
| $k_D$ | $0.1\,s^{-1}$ | MinD membrane attachment rate |
| $k_{dD}$ | $0.1\,\mu m\,s^{-1}$ | MinD membrane recruitment rate |
| $k_{de}$ | $0.5\,s^{-1}$ | MinDE membrane detachment rate |
| $k_{dEr}$ | $2\,\mu m\,s^{-1}$ | Reactive MinE recruitment rate |
| $k_{dEi}$ | $10^{-3}\,\mu m\,s^{-1}$ | Latent MinE recruitment rate |
| $k_{dE}$ | $2\,\mu m\,s^{-1}$ | MinE recruitment rate (skeleton model) |
| $L$ | $500\,\mu m$ | Size of simulation domain |
| $\bar{n}_D$ | $638\,\mu m^{-1}$ | Average MinD density |
| $\bar{n}_E$ | varied | Average MinE density |
| $v_f$ | varied | Flow velocity |

Tab. S1: Parameters used for finite element simulations and linear stability analysis in a line geometry, adopted from Ref.[5]. The rates $\mu$, $k_{dEr}$ and $k_{dEi}$ only apply to the full model with conformational switching of MinE, while the rate $k_{dE}$ describes MinE recruitment in the 'skeleton model'. $\bar{n}_D$ and $\bar{n}_D$ are the spatial averages of total MinD and MinE densities, respectively.

We here state the equations of motion for the switch model, and two simplified versions of it; the reduced switch model, which does not include the cytosolic MinE components, and the skeleton model, where conformational switching of MinE is disabled. For a visualization of the reaction kinetics, refer to main text Fig. 1.

##### 1.4.1 Switch model

The switch model dynamics in 1D (as obtained in Sec. 1.2) read

$$\partial_t m_d = D_m \partial_x^2 m_d + \tilde{f}_d \tag{17a}$$
$$\partial_t m_{de} = D_m \partial_x^2 m_{de} + \tilde{f}_{de} \tag{17b}$$
$$\partial_t c_{DD} + v_f \partial_x c_{DD} = D_c \partial_x^2 c_{DD} + \tilde{f}_{DD} \tag{17c}$$
$$\partial_t c_{DT} + v_f \partial_x c_{DT} = D_c \partial_x^2 c_{DT} + \tilde{f}_{DT} \tag{17d}$$
$$\partial_t c_{Er} + v_f \partial_x c_{Er} = D_c \partial_x^2 c_{Er} + \tilde{f}_{Er} \tag{17e}$$
$$\partial_t c_{Ei} + v_f \partial_x c_{Ei} = D_c \partial_x^2 c_{Ei} + \tilde{f}_{Ei}, \tag{17f}$$

with the reaction terms:

$$\tilde{f}_d = (k_D + k_{dD} m_d) c_{DT} - (k_{dEr} c_{Er} + k_{dEi} c_{Ei}) m_d \tag{18a}$$
$$\tilde{f}_{de} = (k_{dEr} c_{Er} + k_{dEi} c_{Ei}) m_d - k_{de} m_{de} \tag{18b}$$
$$\tilde{f}_{DT} = -(k_D + k_{dD} m_d) c_{DT} + \lambda c_{DD} \tag{18c}$$
$$\tilde{f}_{DD} = k_{de} m_{de} - \lambda c_{DD} \tag{18d}$$
$$\tilde{f}_{Er} = k_{de} m_{de} - k_{dEr} m_d c_{Er} - \mu c_{Er} \tag{18e}$$
$$\tilde{f}_{Ei} = -k_{dEi} m_d c_{Ei} + \mu c_{Er}, \tag{18f}$$

which correspond to the schematic in Fig. 1B in the main text. In addition to the parameters found in Table S1, $k_{dEr} = 2\,\mu m\,s^{-1}$ and $k_{dEi} = 0.001\,\mu m\,s^{-1}$ were used.

In a line geometry, the conservation laws Eq. (7) and Eq. (8) read:

$$\bar{n}_{\mathrm{D}} L = \int_0^L \mathrm{d}x \, (m_{\mathrm{d}} + m_{\mathrm{de}} + c_{\mathrm{DD}} + c_{\mathrm{DT}}) \tag{19}$$

$$\bar{n}_{\mathrm{E}} L = \int_0^L \mathrm{d}x (m_{\mathrm{de}} + c_{\mathrm{Ei}} + c_{\mathrm{Er}}). \tag{20}$$

##### 1.4.2 Reduced switch model

As we have showed in Sec. 1.3, under certain conditions a simplified version of the switch model can be used. The dynamics of this reduced switch model are as follows:

$$\partial_t m_{\mathrm{d}} = D_m \partial_x^2 m_{\mathrm{d}} + \bar{r}_{\mathrm{d}} \tag{21a}$$
$$\partial_t m_{\mathrm{de}} = D_m \partial_x^2 m_{\mathrm{de}} + \bar{r}_{\mathrm{de}} \tag{21b}$$
$$\partial_t c_{\mathrm{DD}} = D_c \partial_x^2 c_{\mathrm{DD}} - v_{\mathrm{f}} \partial_x c_{\mathrm{DD}} + \bar{r}_{\mathrm{DD}} \tag{21c}$$
$$\partial_t c_{\mathrm{DT}} = D_c \partial_x^2 c_{\mathrm{DT}} - v_{\mathrm{f}} \partial_x c_{\mathrm{DT}} + \bar{r}_{\mathrm{DT}}. \tag{21d}$$

Reducing the switch model yields the reaction network displayed in Fig. 1E, stated mathematically:

$$\bar{r}_{\mathrm{d}} = (k_{\mathrm{D}} + k_{\mathrm{dD}} m_{\mathrm{d}}) c_{\mathrm{D}} - \left[ k_{\mathrm{dEi}}^{\mathrm{eff}} + \frac{k_{\mathrm{de}} m_{\mathrm{de}}}{\sqrt{\mu D_c}/k_{\mathrm{dEr}} + m_{\mathrm{d}}} \right] m_{\mathrm{d}} \tag{22a}$$
$$\bar{r}_{\mathrm{de}} = \left[ k_{\mathrm{dEi}}^{\mathrm{eff}} + \frac{k_{\mathrm{de}} m_{\mathrm{de}}}{\sqrt{\mu D_c}/k_{\mathrm{dEr}} + m_{\mathrm{d}}} \right] m_{\mathrm{d}} - k_{\mathrm{de}} m_{\mathrm{de}} \tag{22b}$$
$$\bar{r}_{\mathrm{DT}} = -(k_{\mathrm{D}} + k_{\mathrm{dD}} m_{\mathrm{d}}) c_{\mathrm{DT}} + \lambda c_{\mathrm{DD}} \tag{22c}$$
$$\bar{r}_{\mathrm{DD}} = k_{\mathrm{de}} m_{\mathrm{de}} - \lambda c_{\mathrm{DD}}, \tag{22d}$$

with the effective kinetic rate $k_{\mathrm{dEi}}^{\mathrm{eff}} = k_{\mathrm{dEi}} n_E$.

In the reduced switch model only the total number of MinD proteins

$$\bar{n}_{\mathrm{D}} L = \int_0^L \mathrm{d}x \, (m_{\mathrm{d}} + m_{\mathrm{de}} + c_{\mathrm{DD}} + c_{\mathrm{DT}}) \tag{23}$$

is a conserved quantity, allowing us to omit the total MinE density from the control space analysis.

##### 1.4.3 Skeleton model

The equations for the skeleton model (Fig. 1F) are:

$$\partial_t m_{\mathrm{d}} = D_m \partial_x^2 m_{\mathrm{d}} + \bar{g}_{\mathrm{d}} \tag{24a}$$
$$\partial_t m_{\mathrm{de}} = D_m \partial_x^2 m_{\mathrm{de}} + \bar{g}_{\mathrm{de}} \tag{24b}$$
$$\partial_t c_{\mathrm{DD}} = D_c \partial_x^2 c_{\mathrm{DD}} - v_{\mathrm{f}} \partial_x c_{\mathrm{DD}} + \bar{g}_{\mathrm{DD}} \tag{24c}$$
$$\partial_t c_{\mathrm{DT}} = D_c \partial_x^2 c_{\mathrm{DT}} - v_{\mathrm{f}} \partial_x c_{\mathrm{DT}} + \bar{g}_{\mathrm{DT}} \tag{24d}$$
$$\partial_t c_{\mathrm{E}} = D_c \partial_x^2 c_{\mathrm{E}} - v_{\mathrm{f}} \partial_x c_{\mathrm{E}} + \bar{g}_{\mathrm{E}}, \tag{24e}$$

with the reaction terms

$$\bar{g}_{\mathrm{d}} = (k_{\mathrm{D}} + k_{\mathrm{dD}} m_{\mathrm{d}}) c_{\mathrm{DT}} - k_{\mathrm{dE}} m_{\mathrm{d}} c_{\mathrm{E}} \tag{25a}$$
$$\bar{g}_{\mathrm{de}} = k_{\mathrm{dE}} m_{\mathrm{d}} c_{\mathrm{E}} - k_{\mathrm{de}} m_{\mathrm{de}} \tag{25b}$$
$$\bar{g}_{\mathrm{DT}} = -(k_{\mathrm{D}} + k_{\mathrm{dD}} m_{\mathrm{d}}) c_{\mathrm{DT}} + \lambda c_{\mathrm{DD}} \tag{25c}$$
$$\bar{g}_{\mathrm{DD}} = k_{\mathrm{de}} m_{\mathrm{de}} - \lambda c_{\mathrm{DD}} \tag{25d}$$
$$\bar{g}_{\mathrm{E}} = k_{\mathrm{de}} m_{\mathrm{de}} - k_{\mathrm{dE}} m_{\mathrm{d}} c_{\mathrm{E}}. \tag{25e}$$

Analogously to the switch model, the total MinD and MinE densities

$$\bar{n}_{\mathrm{D}} L = \int_0^L \mathrm{d}x \, (m_{\mathrm{d}} + m_{\mathrm{de}} + c_{\mathrm{DD}} + c_{\mathrm{DT}}) \tag{26}$$

$$\bar{n}_{\mathrm{E}} L = \int_0^L \mathrm{d}x \, (m_{\mathrm{de}} + c_{\mathrm{E}}) \tag{27}$$

are conserved by the above dynamics.

##### 1.5 Details of hysteresis sweeps

To probe the transition between upstream and downstream propagation in the phase diagram in Fig. 3, we performed sweeps where the flow velocity $v_{\mathrm{f}}$ was varied adiabatically over the course of the simulation.

To study the downstream to upstream transition, we initiated patterns at a flow velocity where patterns propagate downstream, if the simulation is initiated from an initially homogeneous state. After the patterns have stabilized in the typical downstream propagating shape, we linearly increased the flow velocity, with a slope that did not exceed $\mathrm{d}v_{\mathrm{f}}/\mathrm{d}t < 0.03 \mathrm{\mu m\, s^{-2}}$.

Analogously, we studied the upstream to downstream transition starting from an upstream propagating pattern, and decreasing the velocity linearly with a slope smaller (in absolute value) than $|\mathrm{d}v_{\mathrm{f}}/\mathrm{d}t| < 0.03 \mathrm{\mu m\, s^{-2}}$.

To completely map out the bistability regime, we also performed simulations where the pattern was initiated at a point in parameter space which exhibits exclusively downstream propagation and then increased the total MinE density, by uniformly supplying it across the entire domain. The transitions from downstream to upstream propagation obtained in this manner are indicated by upright triangles in panel A of Fig. S3. The rate at which the density was changed was $\mathrm{d}n_E/\mathrm{d}t < 0.5 \mathrm{\mu m^{-1}\, s^{-1}}$.

##### 1.6 Details of simulations with disabled MinE advection

To unravel the mechanism behind downstream propagation at low E:D ratios, we performed simulations where we prohibited advection of the cytosolic MinE species by setting $v_{\mathrm{f}} = 0$ in the equations of motion for the species $c_{\mathrm{Er}}$ and $c_{\mathrm{Ei}}$. Strikingly, the patterns still propagated downstream – for typical profile shapes, side by side with typical profile shapes where all species advect normally see Fig. S5.

##### 1.7 Linear Stability Analysis

To study the dynamics in the vicinity of the homogeneous steady state, we performed linear stability analysis (LSA). Collecting the concentrations into a vector $\mathbf{u}(x,t) = [u_1(x,t), ..., u_n(x,t)]^T$, we can write a general Reaction-Diffusion-Advection equation as

$$\partial_t \mathbf{u} = \underline{\underline{D}} \partial_x^2 \mathbf{u} - \underline{\underline{V}} \partial_x \mathbf{u} + \mathbf{f}(\mathbf{u}), \tag{28}$$

where $\underline{\underline{D}} = \mathrm{diag}(D_1, ..., D_n)$ contains the diffusion constants.
The velocity matrix $\underline{\underline{V}} = \mathrm{diag}(v_{\mathrm{f}}, 0, ..., v_{\mathrm{f}})$ determines whether or not a certain component is advected or not (i.e. it is set to zero for membrane concentrations and $v_{\mathrm{f}}$ for cytosolic components). The vector of nonlinear functions $\mathbf{f} = [f_1(\mathbf{u}), ..., f_n(\mathbf{u})]^T$ encodes the reaction kinetics.

We consider the dynamics of small perturbations $\delta \mathbf{u}$ to the homogeneous steady state $\mathbf{u}^*$, for which $\mathbf{f}(\mathbf{u}^*) = \mathbf{0}$. Inserting $\mathbf{u} = \mathbf{u}^* + \delta \mathbf{u}$ into Eq. (28) and linearizing the reaction kinetics, we obtain:

$$\partial_t \delta \mathbf{u} = \underline{\underline{D}} \partial_x^2 \delta \mathbf{u} - \underline{\underline{V}} \partial_x \delta \mathbf{u} + \underline{\underline{J}} \delta \mathbf{u}, \tag{29}$$

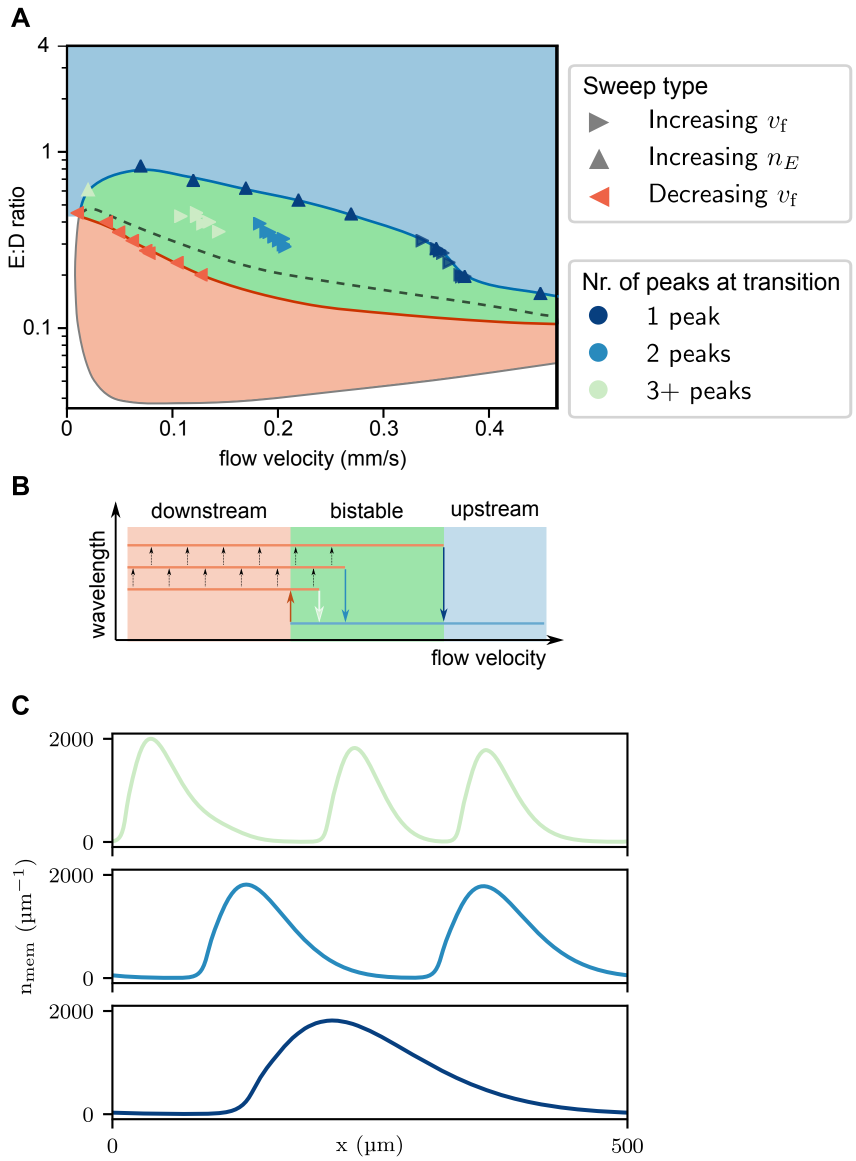


Fig. S3: **A** Phase diagram of the full model, where the shadings correspond to the downstream, bistable and upstream regime, for red, green and blue, respectively (same as in Fig. 3A). The dashed line marks the E:D ratios above which upstream propagating modes are unstable in a Linear Stability Analysis (see also the corresponding dashed line and caption in Fig. S6, panel A). The markers correspond to where reversals of the propagation direction happened in sweeps, where a parameter was varied adiabatically (see SI Sec. 1.5). For increasing $v_\mathrm{f}$ and increasing $n_E$ the transition was from downstream to upstream, while the decreasing $v_\mathrm{f}$ correspond to upstream to downstream transitions. The downstream to upstream markers are color coded to display the wavelength of the pattern at the transition point – we quantify that by the number of peaks (see panel C). **B** Similar to main text Fig. 3B, but with the transition arrows color coded in agreement with panels A and C. **C** Different wavelength pattern profiles of the downstream propagating patterns shortly before the transition point.

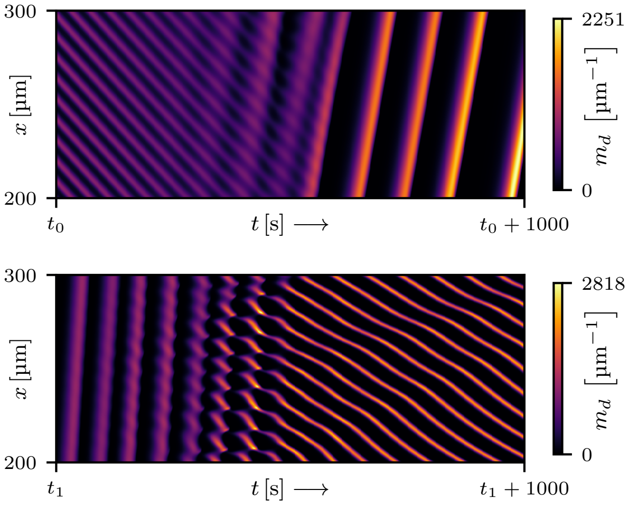


Fig. S4: Typical kymographs of the upstream to downstream (above) and downstream to upstream transition (below). Advective flow runs in the positive $x$-direction.

where $\left(\underline{\underline{J}}\right)_{ij} = \frac{\partial f_i}{\partial u_j}|_{\mathbf{u}^*}$.

Using the ansatz

$$\delta \mathbf{u}^{(q)}(x,t) = e^{\sigma t + iqx}\mathbf{e}^{(q)} \tag{30}$$

Eq. (29) turns into an eigenvalue problem

$$-q^2\underline{\underline{D}}\mathbf{e}^{(q)} - iq\underline{\underline{V}}\mathbf{e}^{(q)} + \underline{\underline{J}}\mathbf{e}^{(q)} = \sigma(q)\mathbf{e}^{(q)}, \tag{31}$$

with $\mathrm{Re}\left[\sigma(q)\right]$ representing the growth rate of a perturbation with a wavenumber $q$. In addition, due to the travelling wave form of the ansatz in Eq. (30), we recognize in $-\mathrm{Im}\left[\sigma(q)\right]$ the angular frequency $\omega(q)$, which allows us to extract the LSA expression for the phase velocity[13] as

$$v_p(q) = \frac{-\mathrm{Im}\left[\sigma(q)\right]}{q}. \tag{32}$$

As $\mathrm{Im}\left[\sigma(q)\right]$ typically changes sign within the band of the unstable modes $q$, it is a priori unclear which (if any) of the wavenumbers $q$ would be relevant to determine the propagation direction. It turns out that in general, the sign of the imaginary part across the band of the unstable nodes is not a good proxy for the actual propagation direction observed in numerical simulations. For the parameter set used throughout this manuscript however, we obtain good agreement between the LSA prediction and the direction of propagation (see panel A in Fig. S6). We believe that this is a consequence of the separation of the typical length scales of the mechanisms underlying the downstream and upstream propagating instabilities – see the bimodal structure of the dispersion relations in panel B of Fig. S6. This bimodal structure of the dispersion relations further corroborates the bistability observed in numerical simulations and experiments.

##### 1.8 Other models of Min protein dynamics

We performed a preliminary analysis of how flow influences pattern formation in some of the other models for the Min protein dynamics in the literature. In the following, we provide the modified part of the dynamics for each of the models we considered, state the parameters used and summarize the observations.

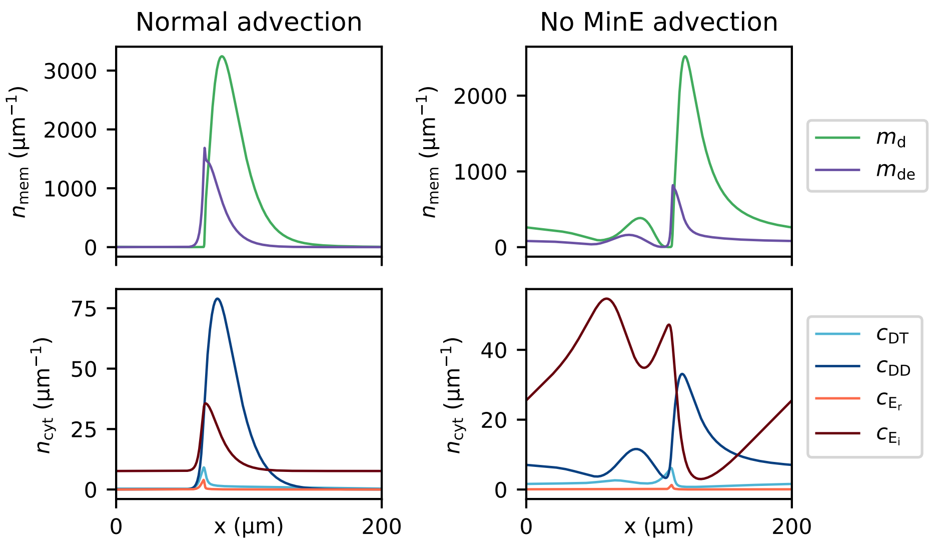


Fig. S5: Typical profile shapes of the downstream propagating patterns, in the case of all cytosolic species being fully advected (left panel), and when exclusively cytosolic MinD is advected (right panel).

##### 1.8.1 Persistent MinE membrane binding

We extended the switch model by persistent MinE membrane binding introduced by Denk, et al.[4] to test whether this has an effect on the propagation direction of waves. The model is essentially an extension of the Switch model, that includes the possibility of MinE persisting on the membrane without being bound in a MinDE complex. We denote this density of membrane bound MinE proteins as $m_e$. The equations of motion are then as for the switch model (see Sec. 1.4.1), with the additional equation governing the dynamics of membrane bound MinE:

$$\partial_t m_e = D_m \partial_x^2 m_e + f_e \tag{33}$$

and the following reaction terms modified or added:

$$f_d = (k_{\mathrm{D}} + k_{\mathrm{dD}} m_{\mathrm{d}}) c_{\mathrm{D}} - (k_{\mathrm{dEr}} c_{\mathrm{Er}} + k_{\mathrm{dEi}} c_{\mathrm{Ei}} + k_{\mathrm{ed}} m_e) m_{\mathrm{d}} \tag{34a}$$
$$f_{de} = (k_{\mathrm{dEr}} c_{\mathrm{Er}} + k_{\mathrm{dEi}} c_{\mathrm{Ei}} + k_{ed} m_e) m_{\mathrm{d}} - k_{de} m_{\mathrm{de}} \tag{34b}$$
$$f_e = k_{de} m_{\mathrm{de}} - k_{ed} m_e m_{\mathrm{d}} - k_e m_e \tag{34c}$$
$$f_{Er} = k_e m_e - k_{\mathrm{dEr}} m_{\mathrm{d}} c_{\mathrm{Er}}. \tag{34d}$$

For the parameter range studied in Ref.[4], we found no qualitative change compared to the model without MinE membrane binding, i.e. what we here refer to as the switch model.

##### 1.8.2 Loose, et al. model

Next, we performed simulations of the model introduced by Loose, et al.[8] extended to include the MinE conformational switch. This amounts to modifying the reactions given in the original

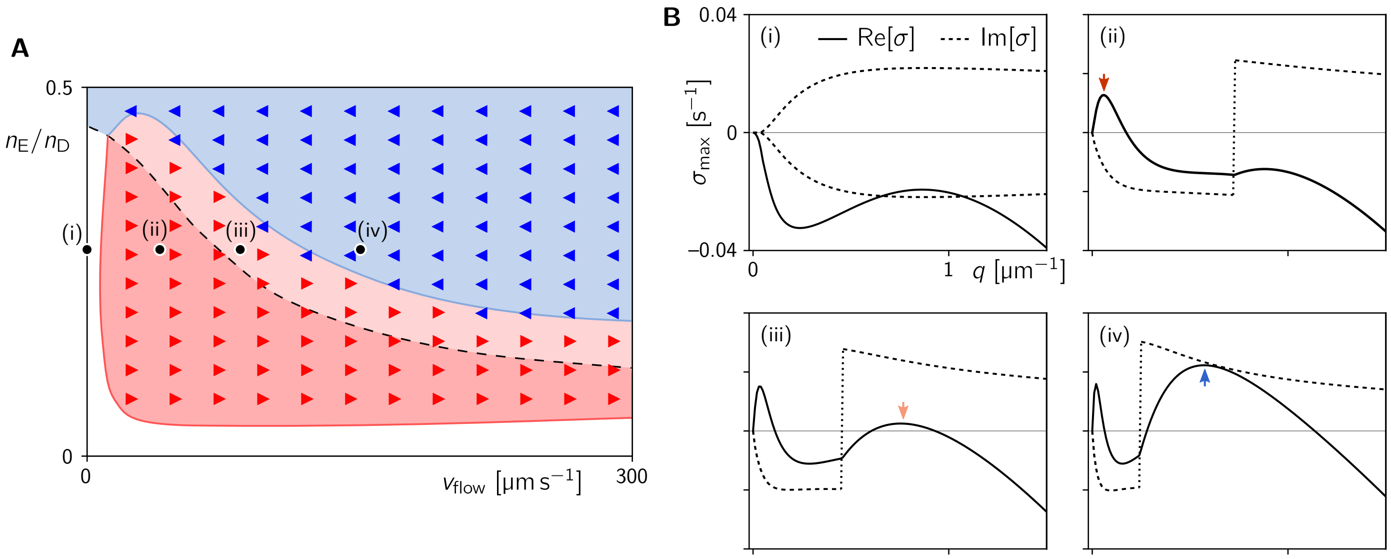


Fig. S6: **A** Stability diagram obtained from linear stability analysis (background shading) and numerical simulations (triangle markers) of the full model in 1D. The shading is based on the signs of the imaginary part at the fastest growing mode $q_{\mathrm{max}}$ and at the right edge of the band of unstable modes $q_+$ (see representative dispersion relations in B and text for details). Blue and red markers indicate upstream and downstream propagation observed in numerical simulations initiated at the homogeneous steady state. **B** Dispersion relations corresponding to the points marked in the phase diagram (A). In the absence of flow [panel (i)], the eigenvalues come in pairs of complex conjugates, such that there are two branches with equal real part (solid line) and opposite imaginary parts (dashed lines). This reflects the left-right symmetry of the system. This symmetry, and thus the degeneracy of the real parts is broken by flow. In panels (ii–iv), only the eigenvalue with the largest real part is plotted causing the jumps in the imaginary part.

reference as:

$$
\begin{aligned}
f_d &= (\omega_D + \omega_{dD} m_{\mathrm{d}}) c_{\mathrm{D}} - m_{\mathrm{d}} \big[ c_{\mathrm{Ei}} (\omega_{Ei} + \omega_{eEi} m_{\mathrm{de}}^2) \\
&\quad + c_{\mathrm{Er}} (\omega_{Er} + \omega_{eEr} m_{\mathrm{de}}^2) \big] && \text{(35a)}\\
f_{de} &= m_{\mathrm{d}} \left[ c_{\mathrm{Ei}} (\omega_{Ei} + \omega_{eEi} m_{\mathrm{de}}^2) + c_{\mathrm{Er}} (\omega_{Er} + \omega_{eEr} m_{\mathrm{de}}^2) \right] - \omega_{de} m_{\mathrm{de}} && \text{(35b)}\\
f_D &= \omega_{de} m_{\mathrm{de}} - (\omega_D + \omega_{dD} m_{\mathrm{d}}) c_{\mathrm{D}} && \text{(35c)}\\
f_{Ei} &= -c_{\mathrm{Ei}} m_{\mathrm{d}} (\omega_{Ei} + \omega_{eEi} m_{\mathrm{de}}^2) + \mu c_{\mathrm{Er}} && \text{(35d)}\\
f_{Er} &= \omega_{de} m_{\mathrm{de}} - c_{\mathrm{Er}} m_{\mathrm{d}} (\omega_{Er} + \omega_{eEr} m_{\mathrm{de}}^2) - \mu c_{\mathrm{Er}}. && \text{(35e)}
\end{aligned}
$$

All of the reaction rates and diffusion constants were taken from the original publication[8], supplemented with $\mu = 100\,\mathrm{s}^{-1}$, $\omega_{eEi} = 1 \cdot 10^{-20}\,\mathrm{\mu m^2\,s^{-1}}$, $\omega_{eEr} = 2.1 \cdot 10^{-18}\,\mathrm{\mu m^2\,s^{-1}}$, $\omega_{Ei} = 9.5 \cdot 10^{-11}\,\mathrm{\mu m\,s^{-1}}$, $\omega_{Er} = 1.9 \cdot 10^{-8}\,\mathrm{\mu m\,s^{-1}}$.

We simulated this model on an interval of length $L = 500\,\mathrm{\mu m}$, and observed exclusively downstream propagation both with and without MinE molecular switching, for several densities in the range $n_E = 2.5 \cdot 10^6 - 1.9 \cdot 10^7\,\mathrm{\mu m^{-1}}$. We did however notice that the regime pattern formation is extended to higher E:D ratios upon the introduction of the MinE conformational switch.

##### 1.8.3 Bonny, et al. model

Finally, we performed a preliminary analysis of the effects of flow on the model introduced by Bonny, et al.[1]. As this model was introduced in the 2+1D slice geometry as introduced in

Sec. 1.1, we reuse that framework with the following reaction kinetics, which constitute our extension of the reaction kinetics given in the original reference:

$$
\begin{aligned}
f_d &= (\omega_D + \omega_{dD} m_\mathrm{d}) c_\mathrm{D} (c_\mathrm{max} - m_\mathrm{d} - m_\mathrm{de})/c_\mathrm{max} \\
&\quad - (\omega_{Er} c_\mathrm{Er} + \omega_{Ei} c_\mathrm{Ei} + \omega_{ed} m_e) m_\mathrm{d} && \text{(36a)} \\
f_{de} &= (\omega_{Er} c_\mathrm{Er} + \omega_{Ei} c_\mathrm{Ei} + \omega_{ed} m_e) m_\mathrm{d} - (\omega_{de,m} + \omega_{de,c}) m_\mathrm{de} && \text{(36b)} \\
f_e &= \omega_{de,m} m_\mathrm{de} - \omega_{ed} m_e m_\mathrm{d} - \omega_e m_e && \text{(36c)} \\
f_D &= -(\omega_D + \omega_{dD} m_\mathrm{d}) c_\mathrm{D} (c_\mathrm{max} - m_\mathrm{d} - m_\mathrm{de})/c_\mathrm{max} \\
&\quad + (\omega_{de,m} + \omega_{de,c}) m_\mathrm{de} && \text{(36d)} \\
f_{Er} &= \omega_e m_e + \omega_{de,c} m_\mathrm{de} - \omega_{Er} m_\mathrm{d} c_\mathrm{Er} && \text{(36e)} \\
f_{Ei} &= -\omega_{Ei} m_\mathrm{d} c_\mathrm{Ei}. && \text{(36f)}
\end{aligned}
$$

We note here explicitly, that the switching of MinE from its reactive to its latent conformation was modelled in the same way as in the switch model, using a linear process with a rate $\mu = 100\,\mathrm{s}^{-1}$. However, the Bonny model does not include the binding of ATP to MinD explicitly, i.e. there is only a single MinD species in their model. Another addition to the switch model discussed in Sec. 1.4.1 is the inclusion of membrane bound MinE, which is governed by Eq. (33) with $D_m$ replaced by the original $D_e$. All of the parameters were taken from the original paper, with the exception of $\omega_{Ei} = 1.36 \cdot 10^{-5}\,\mathrm{\mu m^2\,s^{-1}}$, $\omega_{Er} = 1.36 \cdot 10^{-2}\,\mathrm{\mu m\,s^{-1}}$ and the total MinE density which was varied in the range $n_\mathrm{E} = 2.5 \cdot 10^6\,\mathrm{\mu m^{-1}} - 1.9 \cdot 10^7\,\mathrm{\mu m^{-1}}$.

Upon introducing MinE molecular switching in this manner, we observed that the range of pattern formation was extended to higher E:D ratios. However, the propagation direction was downstream both with and without MinE switching, in the entirety of the pattern forming range.

#### 2 Experimental Methods (extended)

##### 2.1 Chemicals

Unless indicated otherwise, chemicals were bought from Sigma Aldrich. For buffer and solvent preparation, we exclusively used deionized water (Merck Mili-Q®). An overview on important reagents and their storage conditions is given in Tab. S2.

| Product name | Abbr. | Provider | Article number | Storage |
|---|---|---|---|---|
| 18:1 (Δ9-Cis) PC | DOPC | Avanti Polar Lipids | 850375C-25mg | chloroform, −20 °C |
| 18:1 (Δ9-Cis) PG | DOPG | Avanti Polar Lipids | 840475C-25mg | chloroform, −20 °C |
| TopFluor® Cardiolipin | - | Avanti Polar Lipids | 810286C-100ug | chloroform, −20 °C |
| Adenosine triphosphate | ATP | Thermo Fisher | 10304340 | −20 °C |
| Pyruvate kinase | PK | Merck Sigma | P7768-1KU | stock at 4 °C, aliquots at −20 °C |
| Phospho-enolpyruvic acid monopotassium salt, 99 % | PEP | Alfa Aesar | B20358.06 | dissolved in water and pH adjusted with KOH to ∼7.4, stored at −20 °C |

Tab. S2: Overview on important reagents and their storage conditions

##### 2.2 Preparation and handling of SUVs

Chloroform-suspended lipids (DOPG:DOPC in a ratio 33:67, with 0.01–0.02 % mol TopFluor Cardiolipin) were mixed in a glass vial, then dried with a nitrogen gun and further left to dry in a desiccator for 15 minutes. Next, they were resuspended in SUV buffer (150 $mol\,L^{-1}$ KCl, 25 $mol\,L^{-1}$ TRIS pH 7.4-7.5 at room temperature) at a concentration of 5 $mg\,mL^{-1}$ and left to swell at room temperature for at least 1 h. After swelling, they were vortexed for about 20 s and sonicated for 30 min. Avanti Mini-Extruder (Avanti Polar Lipids) was used to produce SUVs. The extruder was pre-warmed on a hotplate set to 45 °C and remained there throughout the procedure. The lipid solution was extruded through a filter membrane of pore size 100 nm for 21 times, followed by another round of extrusion using filters with a pore size of 40 nm. The uneven number of filtering counts ensured that the lipid solution ended up in the originally empty one of the two syringes, so as to avoid taking along aggregates that might accumulate in the first syringe. The resulting SUV solution was aliquoted and stored at 4 °C for up to 10 days. Before the experiment, an aliquot of SUV solution (stored at 5 $mg\,mL^{-1}$ at 4 °C) was diluted to a concentration of 1 $mg\,mL^{-1}$ in 150 $mmol\,L^{-1}$ KCl, 25 $mmol\,L^{-1}$ TRIS (pH 7.4-7.5) and 4 $mmol\,L^{-1}$ $MgCl_2$, then sonicated for 15 min. Next, the solution was injected into the flow cell and incubated at 37 °C for 1 h. The flow cell was then rinsed with Min buffer (150 $mmol\,L^{-1}$ KCl, 25 $mmol\,L^{-1}$ TRIS pH 7.4-7.5 and 5 $mmol\,L^{-1}$ $MgCl_2$) using at least 10 times the volume of the flow cell to ensure removal of residual SUVs.

##### 2.3 Protein purification and labeling

Min protein were purified and labeled as described previously (see Caspi & Dekker [3]) and stored in Min storage buffer (containing 50 mmol L$^{-1}$ Hepes pH 7.25 at 4 °C, 150 mmol L$^{-1}$ KCl, 10 % V/V glycerol, 0.1 mmol L$^{-1}$ EDTA pH 7.4 and for MinD additionally 80 mmol L$^{-1}$ ADP) at −80 °C. labeled protein had labeling efficiencies of about 88 % for Cy3:MinD-Lysine and 45 % for Cy5:MinE-Cysteine.

MinE-L3E/I24N (see Denk et al. [5]) was overexpressed, purified and labeled with Cy5-maleimide (Cytiva PA15131) as described previously (see Caspi & Dekker [3]) with the following modifications: Affinity purified proteins were concentrated using a Vivaspin 10 kDa ultrafiltration device before application to a Superdex S200 gel filtration column. The labeling efficiency was about 58 %.

##### 2.4 Flow cell preparation and assembly

To create inlets and outlets, holes were drilled in cover slides. Microscope slides and cover slides were then placed in Teflon containers and pre-cleaned by sequential sonication in deionized water for 5 min, Acetone for 15 min, KOH (1 mol L$^{-1}$) for 1 h and Ethanol for 1 h. Between sonication steps, the containers were rinsed with deionized water several times to remove the preceding solution. Next, the slides were cleaned with acid Piranha ($H_2SO_4$ and $H_2O_2$ in a ratio 5:1) and after rinsing with plenty of deionized water, stored in fresh Ethanol.

To assemble a flow cell, a set of up to four channels was cut into a double layer of Parafilm® using a cutting mold. Microscope and cover slides were dried with a nitrogen gun, then additionally plasma-cleaned (at about 50 W for 1 min) before assembly. Cover slide, parafilm and microscope slide were stacked on top of each other, then the flow cell was sealed by placing it on a hotplate at 95 °C for a few minutes and gently pressing the channels' edges down with a cotton swab. The flow cells were then stored at room temperature and used either on the same day, or a maximum of two days after assembly. The resulting channels would have heights of 180–230 µm, which were measured on the microscope from the height difference between the top and bottom bilayer. Their cross-sectional widths would be roughly 3 mm and also measured directly on the microscope stage for each individual flow channel. Measurements are given in Tab. S3. As found previously by Brauns et al., coupling of patterns on the upper and lower glass slide can be expected for bulk heights up to 30 µm [2]. To avoid possible complications, it was made sure that all used flow cells considerably exceeded this height.

Cut pipette tips were used to connect the flow channel's inlet and outlet to the tubing, and fixed in position using 5-Minute Epoxy (Thorlabs). For closed-circle experiments, both the channel and the tubing were filled with protein solution prior to connecting and sealing off them with 5-Minute Epoxy. The tubing would go through a dispensing pump (model IPC, Ismatec®), allowing for cycling of the protein solution within a closed system (compare Fig. S7). This system was used for all shown experiments performed with MinE-wiltype.

Due to protein sticking and resulting change in concentration, the closed-circle system was found unsuitable for experiments with the less robust mutant MinE-L3I24N. For experiments with this variant, we used an open system. Here, only the outlet was connected to the pump, while the inlet's tubing led to a reservoir of Min protein solution, from which it could be sucked into the channel.

##### 2.5 Min pattern acquisition

Min protein at the indicated concentrations (with 10–20 % labeled protein, not accounting for labeling efficiency) were supplemented with 0.01 mg mL$^{-1}$ PK and 5 mol L$^{-1}$ PEP and mixed in Min buffer. The volume of all reagents was accounted for to achieve a total concentration of 150 mmol L$^{-1}$ KCl due to its strong influence on Min pattern formation, as found by Vecchiarelli

et al.[12]. The exact final concentration of all buffer components (taking the storage buffer into account) would then amount to 150 mmol L$^{-1}$ KCl, 5 mmol L$^{-1}$ $MgCl_2$, 19–23 mmol L$^{-1}$ TRIS pH 7.4-7.5 (21 °C) and 5–11 mmol L$^{-1}$ Hepes pH 7.25 (4 °C), plus small amounts of glycerol, EDTA and ADP. Pipetting and mixing of Min protein was done using low-binding tubes and pipette tips (Eppendorf™) to reduce the loss of protein while transferring.
Min patterns were incubated for at least 1 h at room temperature before imaging. All images were captured on an Olympus IX-81 inverted microscope with an Andor Revolution XD spinning disk system and FRAPPA, EM-CCD Andor iXon X3 DU897 camera, Andor Revolution and Yokogawa CSU X1 system for illumination and detection, a z-piezo stage and motorized x-y stage, using a 20× objective (Olympus PlanN 20x / 0.4 NA). For excitation of MinD-Cy3 and MinE-Cy5, laser lines 561 nm (617/73 nm filter) and 640 nm (685/40 nm filter) were used. Images were acquired in the central third of the channel (dashed rectangle in Fig. S7) in multiple positions at 15 s intervals (4 frames per minute). Acquisition time was set to 400 ms, laser power was adjusted individually depending on the protein concentration.

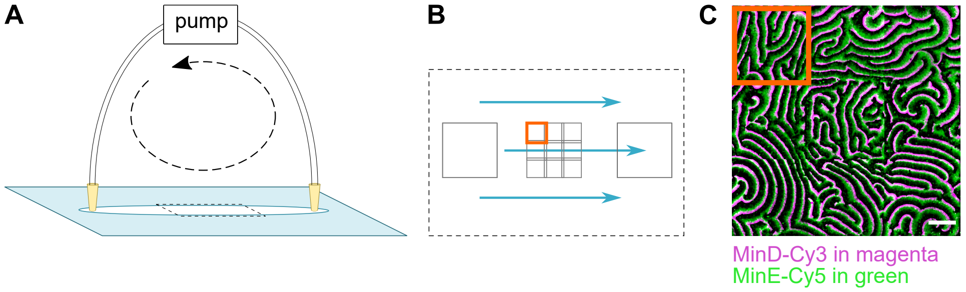


Fig. S7: **A** Closed-circle flow cell setup (schematic). **B** Stitching of up to three 3x3 regions, reaching areas of about 0.8 mm x 0.8 mm in size. Dashed rectangle corresponds to region indicated in A. **C** Exemplary stitched image frame, stitched from three frames of originally 512x512 pixel. Scale bar 100 µm. Single FOV highlighted in orange, corresponding to the equally highlighted region in **B**.

### 3 Image and data analysis (extended)

#### 3.1 General information

Min image stack data were pre-processed using home-built MatLab scripts (The Math Works, Inc. MATLAB. Version 2018b, The Math Works, Inc., 2020. www.mathworks.com) and analysed using home-built Python[11] scripts following the methodology described in a separate publication[9]. Schematics and figures were arranged using Inkscape (Inkscape Project, 2020. https://inkscape.org).
In the following, we present an outline of the analysis performed for this paper.

#### 3.2 Image processing

The images for this paper were acquired using a spinning disc confocal microscope. The associated relative high background signal could vary considerably for different E:D ratios (with the concentration of MinE-Cy5 at 45 % labeling efficiency ranging from 0.4–1 µmol L$^{-1}$). In addition, patterns did bleach under prolonged imaging, illumination was typically inhomogeneous and pattern diagnostics per field of view (FOV) could be cumbersome as the length scale of wave patterns is typically large compared to the FOV. Finally, the closed-circle flow cell setup posed additional challenges in the form of objects entering and passing through the FOV, as well as adhering to the surface. As Min proteins might form aggregates over time, more and more of these artefacts would emerge as well as occasionally detach from the surface. To minimize the effects of these experimental limitations, prior to wave pattern analysis we prepared the images as follows:
As we acquired images in 3x3 grids, we first split movies into 9 individual tif-stacks. As in our experimental setup, in these images the flow goes from right to left, we flipped all images horizontally so as to change the flow direction from left to right. This does not change any quantitative or qualitative statements of this paper, as it merely corresponds to changing one's perspective onto the sample from a downwards to a upwards view. First, standard cleaning steps were performed for individual movies:

- Fluorescence bleaching was corrected by normalizing each frame on its mean intensity value.
- An illumination correction map $I_{illum}$ was made by strongly smoothing and averaging all movie images, then normalizing the result to its maximum.
- In addition, a 'static background'-image $I_{stat}$ was made by averaging out all moving (wave pattern) features of the movie stack, leaving only static fluorescent features such as specks, holes and scratches.
- Each movie image was then corrected via the following image operation: $I_{cor} = (I_{movie} - I_{stat})/I_{illum}$.
- Occasionally remaining bright or dark artefacts were removed by outlier cropping.
- Images were smoothed to diminish the effect of remaining sharp edges and artefacts from the cropping.

The cleaned movies were then re-assembled into stitched 3x3 frames. Overlaps between adjoining images were cropped out and residual average intensity levels between individual FOVs were corrected for.
This way, all individual movie stacks were assembled into one movie showing a large area pattern with an optimally visible pattern. In our case, these areas were roughly 0.8 mm x 0.8 mm in size. An overview on the used areas is given in Sec. 4.7.

##### 3.3 Image representation

The microscope images shown in the paper were additionally prepared using Fiji [10]. As acquisitions from different experiments were done at different concentration, they varied in intensity of background and local spikes. Depending on the quality of the individual files, images shown in the figures were additionally processed with background subtraction and/or de-noise methods (outlier removal) in Fiji. Notably, the different treatment of images for purely graphical representation does not affect the quantitative analysis as described below in Sec. 3.4, as the latter is performed independently on image data pre-processed solely in MatLab as described in Sec. 3.2.

##### 3.4 Wave front propagation analysis

As described in Ref. [9], we analyzed local wave speed by performing two steps:
1) identification of wave crests for each frame in a stack, and
2) frame-by-frame comparison of points along these wave crests.
To achieve this, we estimate an optical flow vector field using Horn-Schunck algorithm. [7] From such vector fields, we obtain a phase image for each frame, with phases defined by the wave intensity either going up ('front'), or going down ('wake'). Wave 'crests' were then defined as positions between front and wake, in areas of the upper half of the wave intensity (the latter step excludes analogous wave 'valleys'). For every crest position, we determined the velocity by measurement of the perpendicular crest translation. This translation was obtained by sampling a cross-sectional profile at subpixel resolution in the current and the former image, and measuring the shift in crest peak position. This yielded a list of vectors with components $(v_x, v_y)$ at frame $n$ in position $(x, y)$. Results from individual regions were excluded by application of masks. These regions were identified and saved during the previously performed image cleaning routine and based upon outlier detection, thus aiming to diminish the effect of residual static and mobile features.
From the vector components, an angle $\alpha$ was calculated, which corresponds to the wave crest point's direction of movement with respect to the direction of applied flow (defined as 0°, left to right in processed images). The collected vector components and resulting angles from all frames ($n = 2...20$) were further subjected to statistical analysis.

##### 3.5 Autocorrelation analysis

Complementary to the local wave crest analysis, we also obtained global, image-averaged parameters like spatial wavelength. Spatial autocorrelation analysis was performed on 10 individual images per movie. For each autocorrelation output image, a radial average was recorded, starting from the main central correlation peak. The resulting spatial radial correlation curve was subjected to maxima analysis. The first maximum after radius $R = 0$ indicated the most predominant distance between wave edges, irrespective of propagation direction. This distance was identified as a representative wavelength $\lambda$ for this movie.

### 4 Supplementary data

#### 4.1 Concentration/ratio change in closed-circle settings

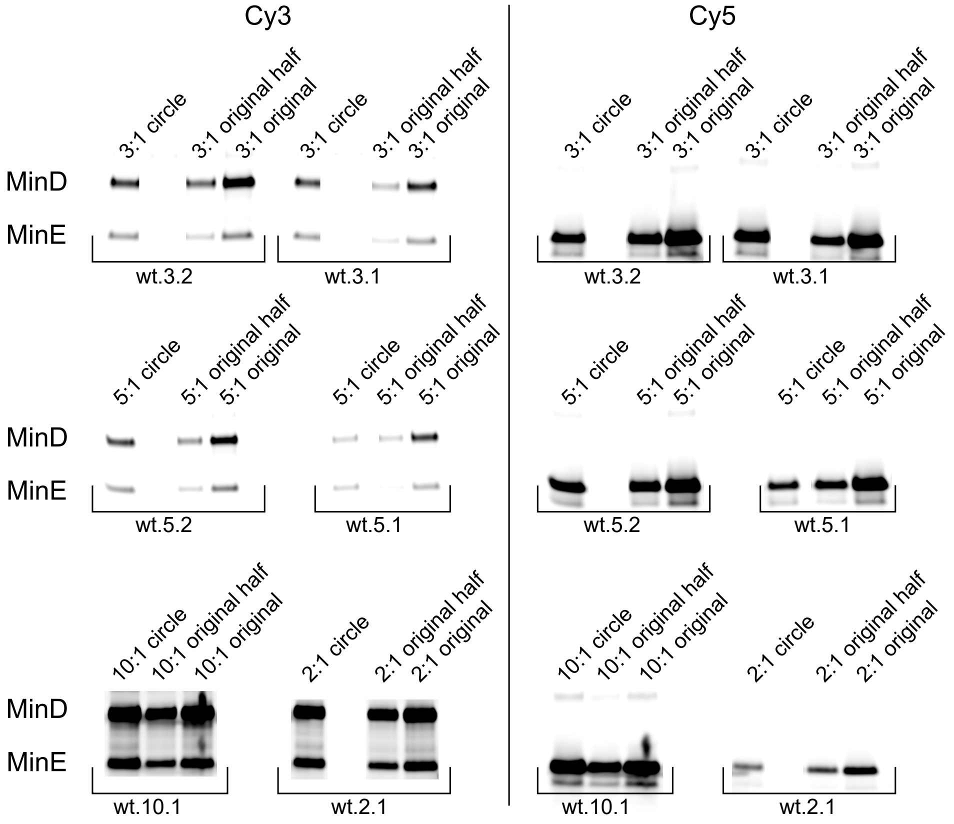


Fig. S8: Results of protein gels run with samples from the experiment IDs indicated, referring to the experiments listed in Tab. S3. For each experiment, a portion of the original sample (*original*), the original sample diluted 1:1 in buffer (*original half*) and the sample extracted from the closed-circle system after the experiment (*circle*) were compared. The remaining fractions of protein were calculated by comparing the intensities of the bands to those of the original sample. The results are summarized in Tab. S3.

Closed-circle experiments (repeated circulation of a fixed amount of Min protein solution thorough a flow channel, led back from the outlet to the inlet via a tubing pump) allows for multiple hours of experimentation without consuming high amounts of protein. However, it can be expected that the concentration of Min protein injected into the flow cell and the tubing differs from the actual concentration in the experimental setting due to sticking of the protein to the inner side of the tubing as well as components of the flow cell not covered by lipid bilayers (e.g. the edge of the parafilm). To estimate this concentration change, the protein solution was ejected from the flow channel and tubing after the experiment and run on a protein gel (NuPAGE™ 4–12 % Bis-Tris-Gel by Initrogen™, in MES buffer) side by side with a portion of the original Min protein solution injected into the channel/tubing. The concentration change was then estimated by comparing the fluorescence signal (Cy3 for MinD, Cy5 for MinE) from both the used and original solution (measured using Amersham Typhoon™, GE Healthcare). The remaining fractions were calculated by interpolation between fluorescence signal of the original solution and the original solution diluted 1:1 in buffer. The gels are shown in Fig. S8, the corrected ratios are shown alongside the initial ones in Tab. S3.
For the sample with initial ratio E:D=10 (*wt.10.1* in Tab. S3), a higher fluorescence was measured for the experiment sample than for the original sample. This was most likely caused by

evaporation or aggregation. Therefore, no correction factor could be calculated for this sample. However, due to the high protein concentration, we assume that the real ratio is close to 10.

##### 4.2 Flow-independent time evolution of wave propagation velocity

As revealed by analysing the wave propagation speed for different flow rates (see Fig. S12), the average propagation speed was found to be lower with flow as compared to without flow. However, due to incubation and acquisition times, a few hours can pass between recording images with and without flow. Therefore, the option has to be considered that the lower propagation speed might be caused by other factors, such as ATP depletion over time. To test this option, two parallel flow channels were filled with Min protein solution ($3\,\mu mol\,L^{-1}$ MinE, $1\,\mu mol\,L^{-1}$ MinD). One was connected to tubing in a closed-circle setup, while the other one was left as it was and checked regularly. The results for the closed-circle experiment are shown in Fig. S9, the results for the control in Fig. S10.

On a side note: Some images such as the one shown in Fig. S9D exhibit heavy bleaching, as can be seen particularly well from dark regions where single microscope images meet. To make sure shortcomings in image quality do not affect quantitative results, such data sets were excluded from analysis based on visual inspection.

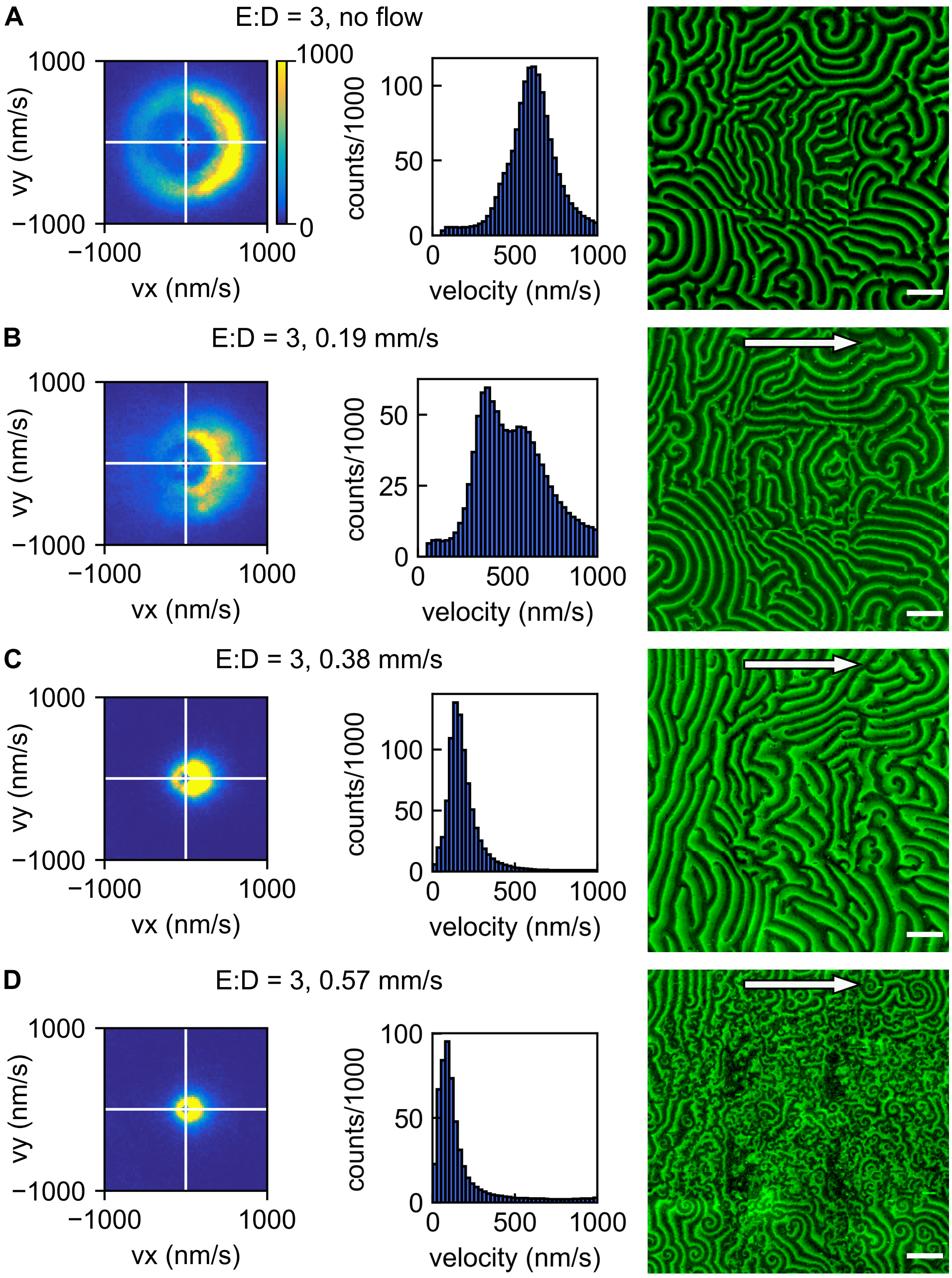


Fig. S9: Initial E:D = 3 (corrected 2.1), wave front propagation analysis performed for MinD-Cy3. Total covered area of 2.01 $mm^2$. Data cropped below 10 % of mean velocity. 2D histogram for segments of 25 $nm\,s^{-1} \times 25\,nm\,s^{-1}$. 1D histograms with bin-size of 25 $nm\,s^{-1}$. Exemplary stitched images for MinE-Cy5 given, scale bars 100 µm. **A** Wave propagation analysis and exemplary image without flow. **B** Wave propagation analysis and exemplary image at 0.19 $mm\,s^{-1}$. **C** Wave propagation analysis and exemplary image at 0.38 $mm\,s^{-1}$. **D** Wave propagation analysis and exemplary image at 0.57 $mm\,s^{-1}$.

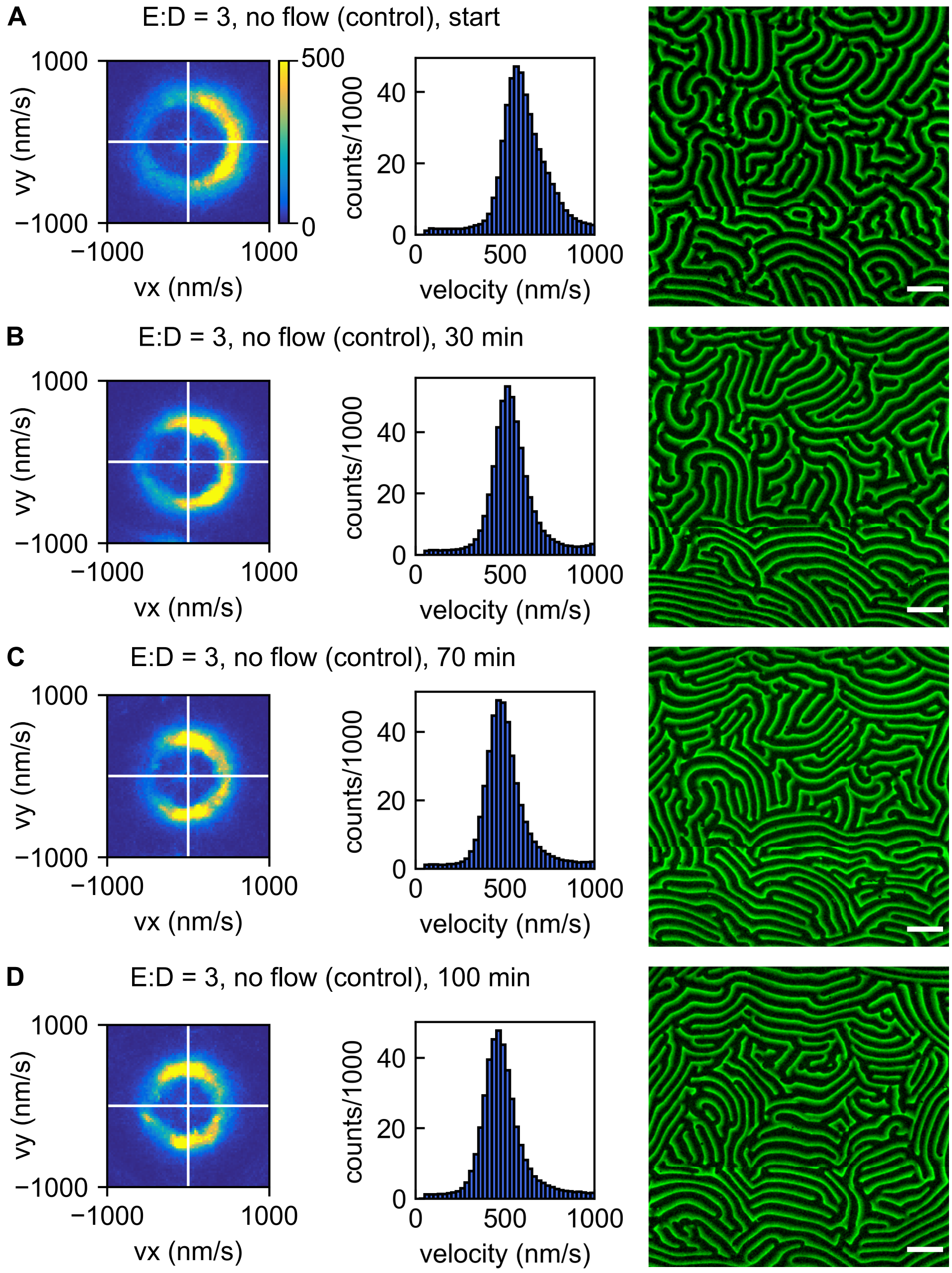


Fig. S10: Initial E:D = 3, wave front propagation analysis performed for MinD-Cy3. Total covered area of 0.69 $mm^2$ (0.83 mm x 0.83 mm). Data cropped below 10 % of mean velocity. 2D histogram for segments of 25 $nm\,s^{-1} \times 25\,nm\,s^{-1}$. 1D histograms with bin-size of 25 $nm\,s^{-1}$. Stitched images for MinE-Cy5 given, scale bars 100 µm. **A** Wave propagation analysis and image at start of experiment. **B** Wave propagation analysis and image about 30 min after start of experiment. **C** Wave propagation analysis and image about 70 min after start of experiment. **D** Wave propagation analysis and image about 100 min after start of experiment.

##### 4.3 Effect of flow reversal

Reversing the flow direction was found to reverse the patterns preferential direction of wave propagation. This was demonstrated during an experiment with 10 µmol L$^{-1}$ MinE, 1 µmol L$^{-1}$ MinD, where the direction of flow was reversed at a flow rate of 0.23 mm s$^{-1}$. See Fig. S11A and Fig. S11B for the results of wave propagation analysis in the initial situations without flow and with flow in the standard direction (left to right). The final patterns shown in in Fig. S11C were acquired about 15 min after flow reversal.

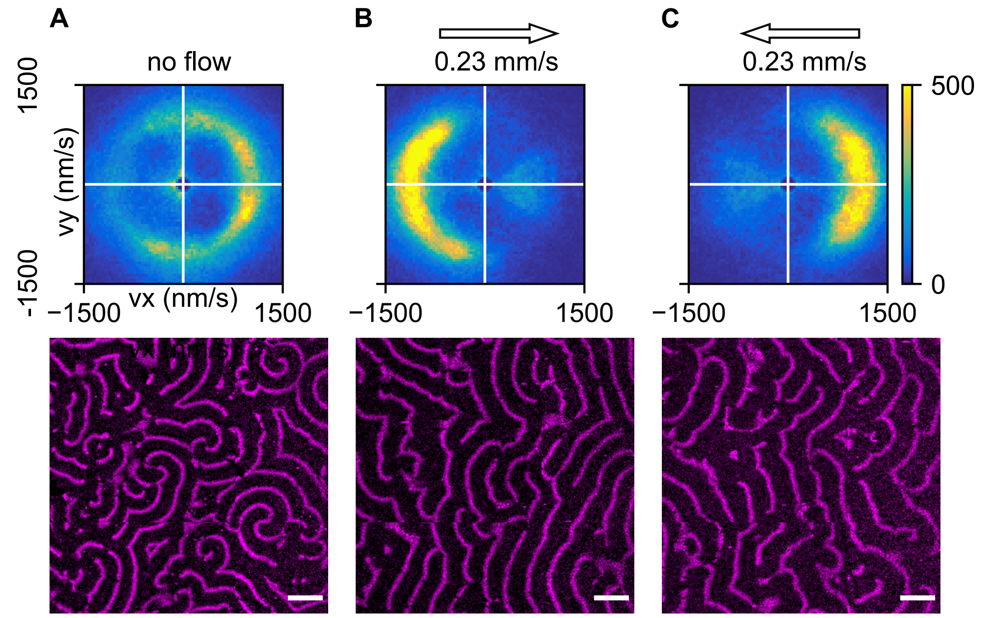


Fig. S11: Initial E:D = 10, wave front propagation analysis performed for MinD-Cy3. The parameter $v_x$ refers to velocity in $x$ (perpendicular to flow), $v_y$ to velocity in $y$ direction (parallel to flow). Shown range of ($v_x$,$v_y$) extends from −1500 to 1500 nm s$^{-1}$. Data cropped below 10 % of mean velocity. 2D histogram for segments of 25 nm s$^{-1}$ × 25 nm s$^{-1}$. Exemplary image shown for MinD-Cy3 channel, scale bar 100 µm. **A** Without flow. **B** With 0.23 mm s$^{-1}$ flow, left to right. **C** With 0.23 mm s$^{-1}$ flow, right to left.

##### 4.4 Wave-crest velocity dependence on flow

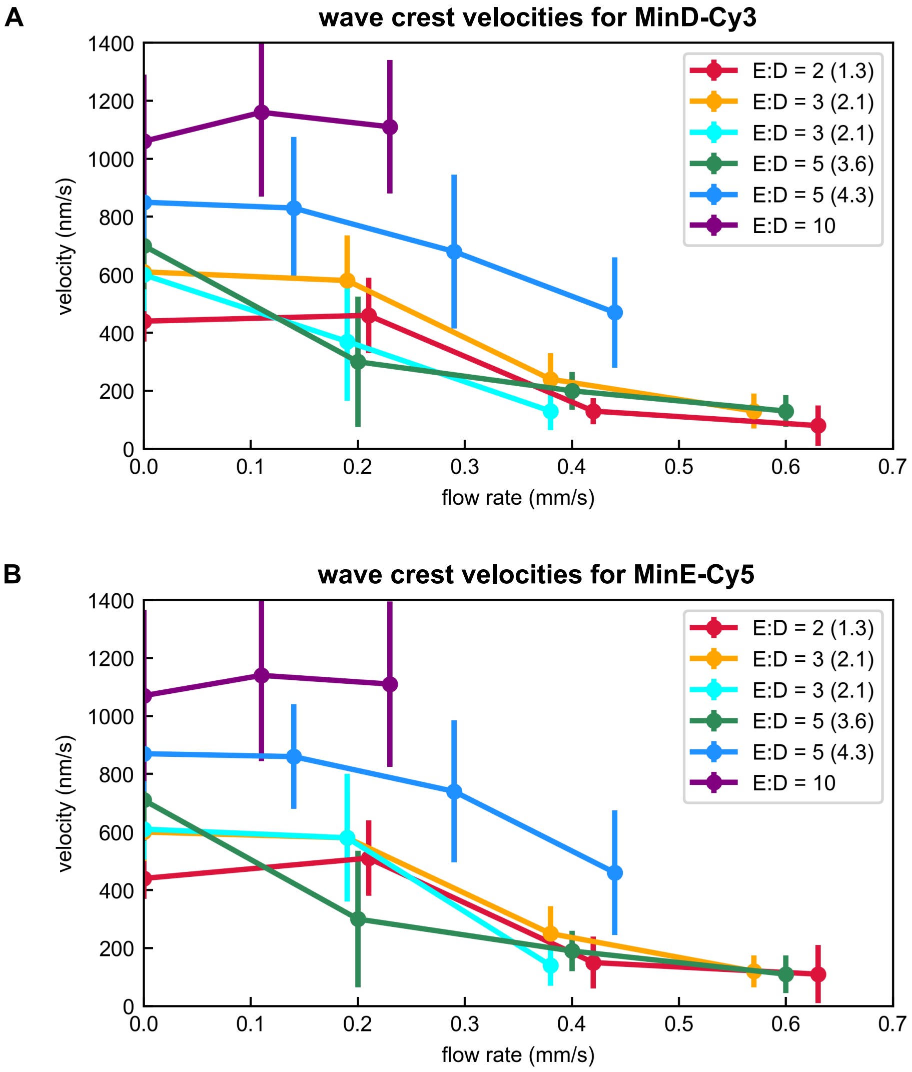


Fig. S12: **A** Dependence of wave velocity for MinD-Cy3 on flow rate and E:D ratio. **B** Dependence of wavelength for MinE-Cy5 on flow rate and E:D ratio. For both **A** and **B**, the wave-crest velocities from all 2-3 imaged regions within one experiment (see Tab. S3) were collected and their magnitude's histogram distribution was calculated with a bining width of $10\,\mathrm{nm\,s^{-1}}$. The peak of this histogram distribution was identified as predominant wave velocity and plotted as a function of the E:D ratio and the flow rate, along with the velocity distribution's FWHM as error bars (peak velocity $\pm$ FWHM/2).

##### 4.5 Wavelength dependence on flow

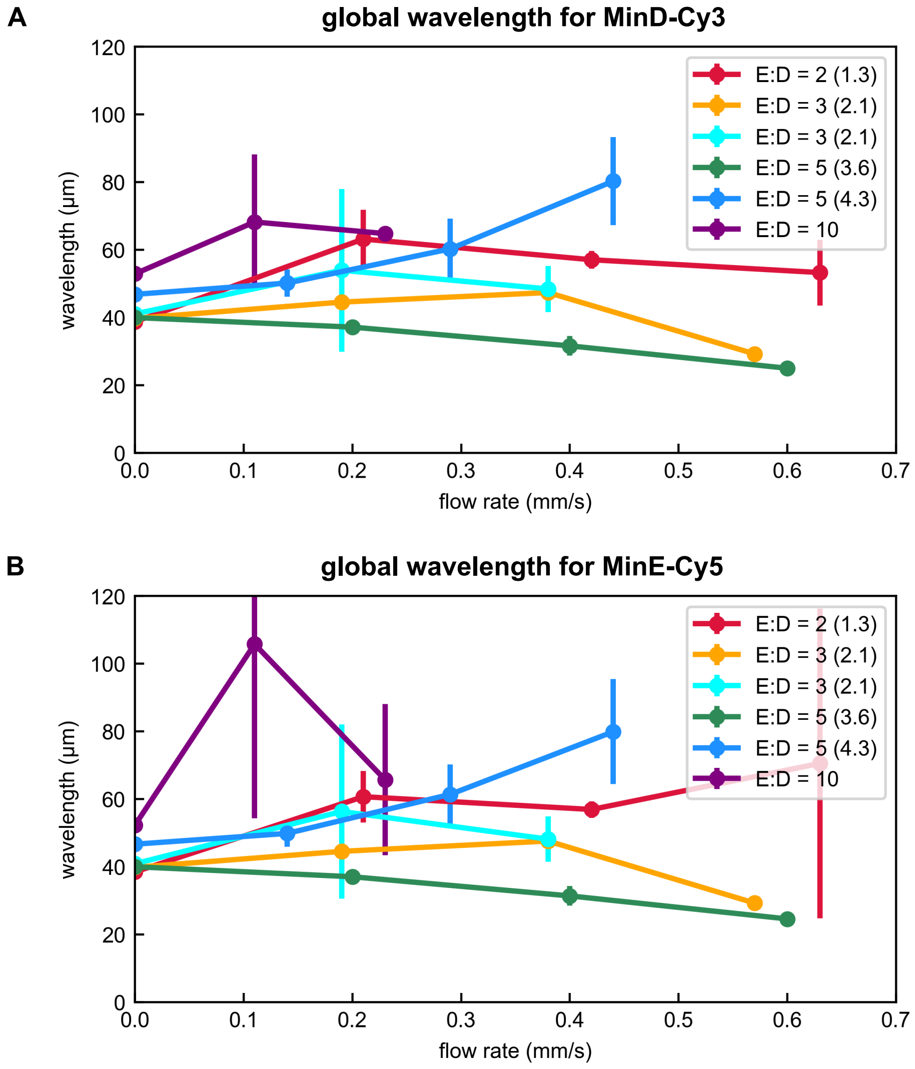


Fig. S13: **A** Dependence of wavelength for MinD-Cy3 on flow rate and E:D ratio. **B** Dependence of wavelength for MinE-Cy5 on flow rate and E:D ratio. For both **A** and **B**, the wavelengths from all 2-3 imaged regions within one experiment (see Tab. S3) were averaged and their standard deviation was calculated to obtain error bars. Large errors could be attributed to issues with image quality, such as bleaching.

##### 4.6 Positional dependence

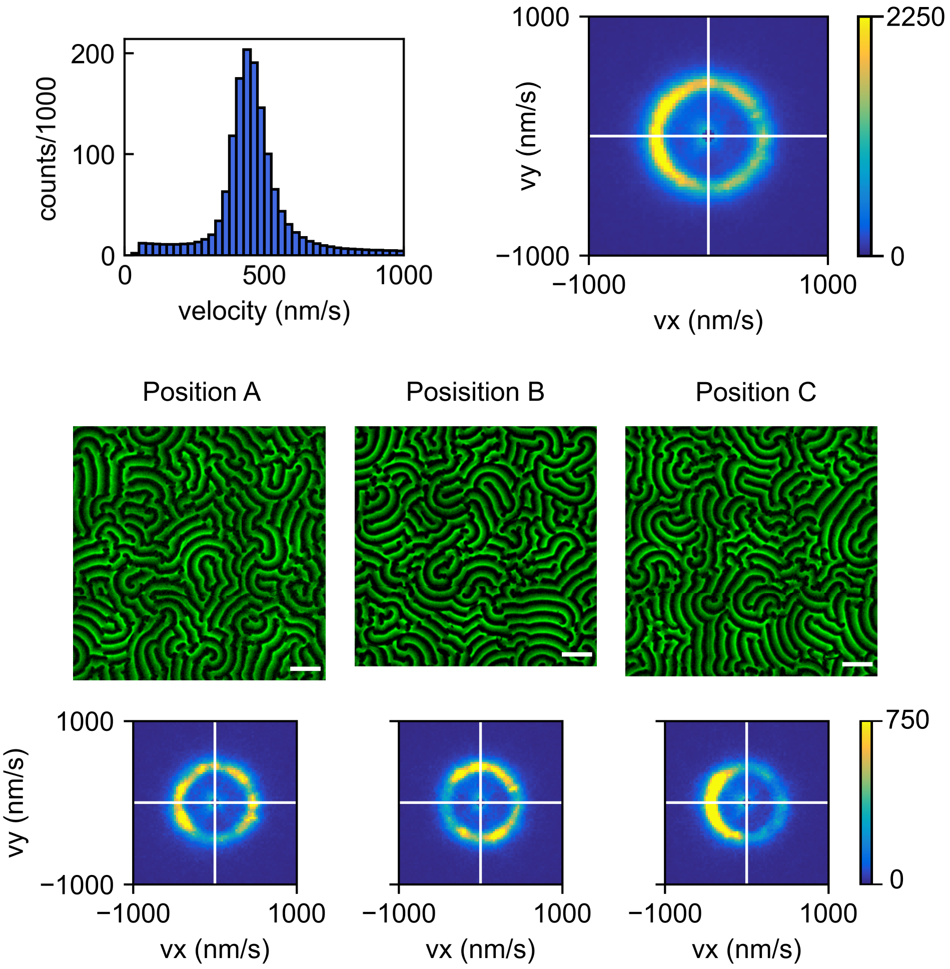


Fig. S14: The wave-crest velocity data for one E:D ratio is obtained from up to three comparable imaging regions within the same flow channel. Here, we show individual analyses of the MinD-Cy3 channel for the three regions imaged at an E:D ratio of 2 (1.3), as fully shown in Fig. S15. Exemplary images shown in green from MinE-Cy5 channel. Scale bars are 100 µm, areas of the three imaging regions are given in Sec. 4.7. The velocity vectors are collected and evaluated with respect to their magnitude and direction. They can be visualized in color-coded 2D histograms (top right and bottom row, in bins of $25\,\mathrm{nm\,s^{-1}} \times 25\,\mathrm{nm\,s^{-1}}$). Their magnitudes can be represented in histogram form (top left, in bins of $25\,\mathrm{nm\,s^{-1}}$).

##### 4.7 Overview on experiments

Tab. S3 on page 28 shows an overview on all image stacks used in this paper. These image stacks are processed as tif files (separate for MinD-Cy3 and MinE-Cy5).
Image IDs starting with "wt" indicate the use of MinE wildtype, IDs starting with "l3" indicate the use of the L3I24N mutant[5].
The flow rate $v$ (always referring to the average cross-sectional flow rate) was calculated from the flow channel's dimensions (height $h$ and width $w$) and the set flow rate $Vt^{-1}$, set on the pump in µL min$^{-1}$, as

$$v = \frac{Vt^{-1}}{w \cdot h}$$

The parameters $v$, $h$ and $w$ allow to calculate the flow channel's Reynolds number. As we obtain low Reynolds numbers for all channels and flow rates, we assume laminar flow for all experiments.
To estimate the standard deviation (error) of the flow rate, the following values were assumed:

- $(Vt^{-1})_{std} = 5\%$ of set value of pump
- $w_{std} = 100\,\mu\text{m}$
- $h_{std} = 10\,\mu\text{m}$

Position 4 of the experiment ID indicates the total area covered by the experiment's type of stitching pattern. The respective values are:

- A: 0.69 mm$^2$ ( 0.83 mm x 0.84 mm )
- B: 0.63 mm$^2$ ( 0.80 mm x 0.79 mm )
- C: 0.68 mm$^2$ ( 0.83 mm x 0.82 mm )
- alt: 0.69 mm$^2$ ( 0.83 mm x 0.83 mm )

For every experiment, the original Min protein mix injected into the flow channel would contain 1 µmol L$^{-1}$ MinD. Therefore, a certain initial ratio corresponds to the same initial concentration of MinE in µmol L$^{-1}$. Both MinD and MinE would partially stick to the channel/tubing. Based on the gels shown in Fig. S8, a corrected ratio was calculated where applicable. Each experiment was performed with a fraction of labeled protein, ranging from 10–20 %, as indicated in Tab. S3. MinD-Cy3 was found to have a labeling efficiency of 88 %, MinE-Cy5 of 45 % (compare Caspi & Dekker[3]).

Tab. S3: Overview on experiments

| image stack ID | initial ratio | corrected ratio | MinD-Cy3 (%) | MinE-Cy5 (%) | width ($\mu$m) | height ($\mu$m) | flow (mm/s) | flow error (mm/s) |
|---|---|---|---|---|---|---|---|---|
| wt.2.1.[A/B/C].00 | 2 | 1.29 | 20 | 20 | 2953 | 183.26 | 0.00 | 0.00 |
| wt.2.1.[A/B/C].01 | 2 | 1.29 | 20 | 20 | 2953 | 183.26 | 0.21 | 0.02 |
| wt.2.1.[A/B/C].02 | 2 | 1.29 | 20 | 20 | 2953 | 183.26 | 0.42 | 0.04 |
| wt.2.1.[A/B/C].03 | 2 | 1.29 | 20 | 20 | 2953 | 183.26 | 0.63 | 0.06 |
| wt.3.1.[A/B/C].00 | 3 | 2.05 | 20 | 20 | 2937 | 203.39 | 0.00 | 0.00 |
| wt.3.1.[A/B/C].01 | 3 | 2.05 | 20 | 20 | 2937 | 203.39 | 0.19 | 0.02 |
| wt.3.1.[A/B/C].02 | 3 | 2.05 | 20 | 20 | 2937 | 203.39 | 0.38 | 0.04 |
| wt.3.1.[A/B/C].03 | 3 | 2.05 | 20 | 20 | 2937 | 203.39 | 0.57 | 0.06 |
| wt.3.2.[A/B/C].00 | 3 | 2.06 | 20 | 20 | 2865 | 208.86 | 0.00 | 0.00 |
| wt.3.2.[A/B/C].01 | 3 | 2.06 | 20 | 20 | 2865 | 208.86 | 0.19 | 0.02 |
| wt.3.2.[A/B/C].02 | 3 | 2.06 | 20 | 20 | 2865 | 208.86 | 0.38 | 0.04 |
| wt.3.2.[A/B/C].03 | 3 | 2.06 | 20 | 20 | 2865 | 208.86 | 0.57 | 0.06 |
| wt.5.1.[A/B/C].00 | 5 | 4.29 | 10 | 10 | 3561 | 229.76 | 0.00 | 0.00 |
| wt.5.1.[A/B/C].01 | 5 | 4.29 | 10 | 10 | 3561 | 229.76 | 0.14 | 0.01 |
| wt.5.1.[A/B/C].02 | 5 | 4.29 | 10 | 10 | 3561 | 229.76 | 0.29 | 0.03 |
| wt.5.1.[A/B/C].03 | 5 | 4.29 | 10 | 10 | 3561 | 229.76 | 0.44 | 0.04 |
| wt.5.2.[A/B/C].00 | 5 | 3.63 | 20 | 20 | 2874 | 198.40 | 0.00 | 0.00 |
| wt.5.2.[A/B/C].01 | 5 | 3.63 | 20 | 20 | 2874 | 198.40 | 0.20 | 0.02 |
| wt.5.2.[A/B/C].02 | 5 | 3.63 | 20 | 20 | 2874 | 198.40 | 0.40 | 0.04 |
| wt.5.2.[A/B/C].03 | 5 | 3.63 | 20 | 20 | 2874 | 198.40 | 0.60 | 0.06 |
| wt.10.1.[A/B].00 | 10 | - | 10 | 10 | 4347 | 230.30 | 0.00 | 0.00 |
| wt.10.1.[A/B].01 | 10 | - | 10 | 10 | 4347 | 230.30 | 0.11 | 0.01 |
| wt.10.1.[A/B].02 | 10 | - | 10 | 10 | 4347 | 230.30 | 0.23 | 0.02 |
| wt.10.1.[A/B].03 | 10 | - | 10 | 10 | 4347 | 230.30 | -0.23 | 0.02 |
| wt.3.3.alt.00 | 3 | - | 20 | 20 | - | - | 0.00 | 0.00 |
| wt.3.3.alt.01 | 3 | - | 20 | 20 | - | - | 0.00 | 0.00 |
| wt.3.3.alt.02 | 3 | - | 20 | 20 | - | - | 0.00 | 0.00 |
| wt.3.3.alt.03 | 3 | - | 20 | 20 | - | - | 0.00 | 0.00 |
| l3.p05.1.[B/C].00 | 0.05 | - | 20 | 10 | 3296 | 218.64 | 0.00 | 0.00 |
| l3.p05.1.[B/C].01 | 0.05 | - | 20 | 10 | 3296 | 218.64 | 0.15 | 0.02 |

#### 4.8 Full pattern direction analysis

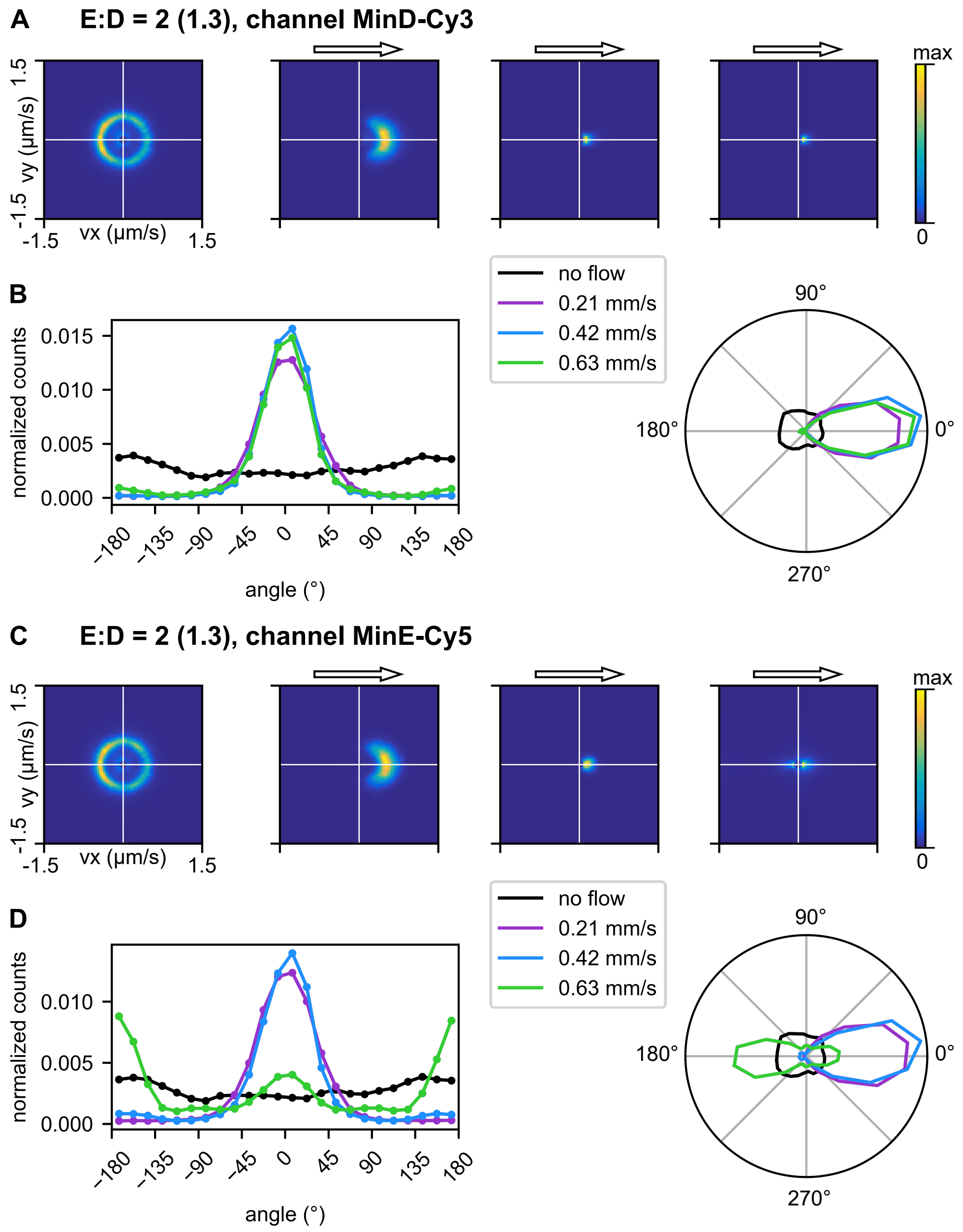


Fig. S15: Analysis for E:D=2 (corrected 1.3), *wt.2.1*, performed with tools presented in Ref.[9] using general smoothing kernel 20 pixels and flow smoothing kernel 50 pixels. Velocity data cropped below 10 % of mean. **A**, **C** Vector counts per segment for no-flow setting and sequentially applied flow rates as given in the box in **B**, represented as 2D histograms (80 bins in $x$-/$y$-direction each), counts per box indicated by color, maximum (yellow) corresponds to maximum bin count for each individual 2D histogram. **B**, **D** 1D histogram and polar histogram representation of counts per direction in bins of 15° for different flow rates.

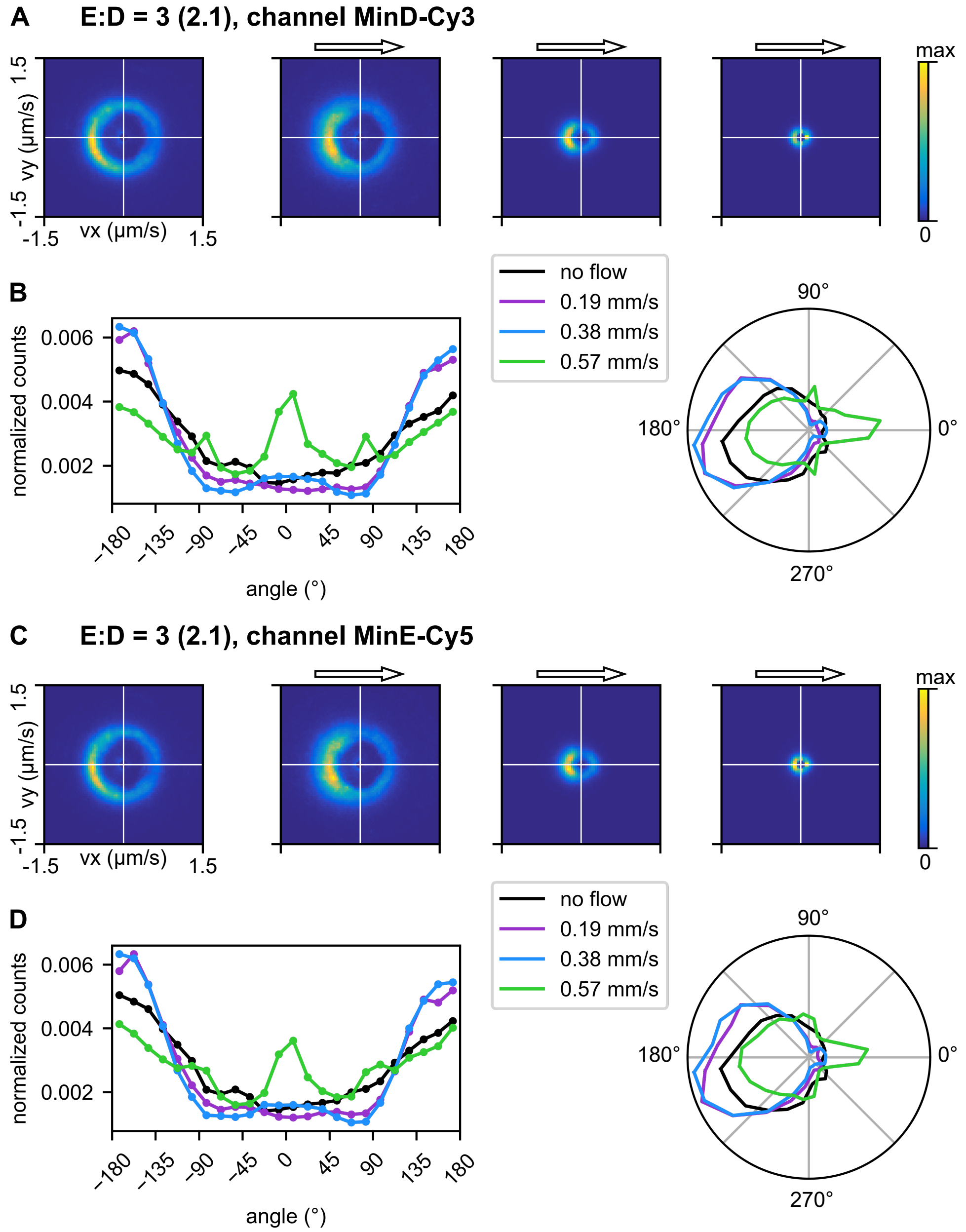


Fig. S16: Analysis for E:D=3 (corrected 2.1), *wt.3.1*, performed with tools presented in Ref.[9] using general smoothing kernel 20 pixels and flow smoothing kernel 50 pixels. Velocity data cropped below 10 % of mean. **A**, **C** Vector counts per segment for no-flow setting and sequentially applied flow rates as given in the box in **B**, represented as 2D histograms (80 bins in $x$-/$y$-direction each), counts per box indicated by color, maximum (yellow) corresponds to maximum bin count for each individual 2D histogram. **B**, **D** 1D histogram and polar histogram representation of counts per direction in bins of 15° for different flow rates.

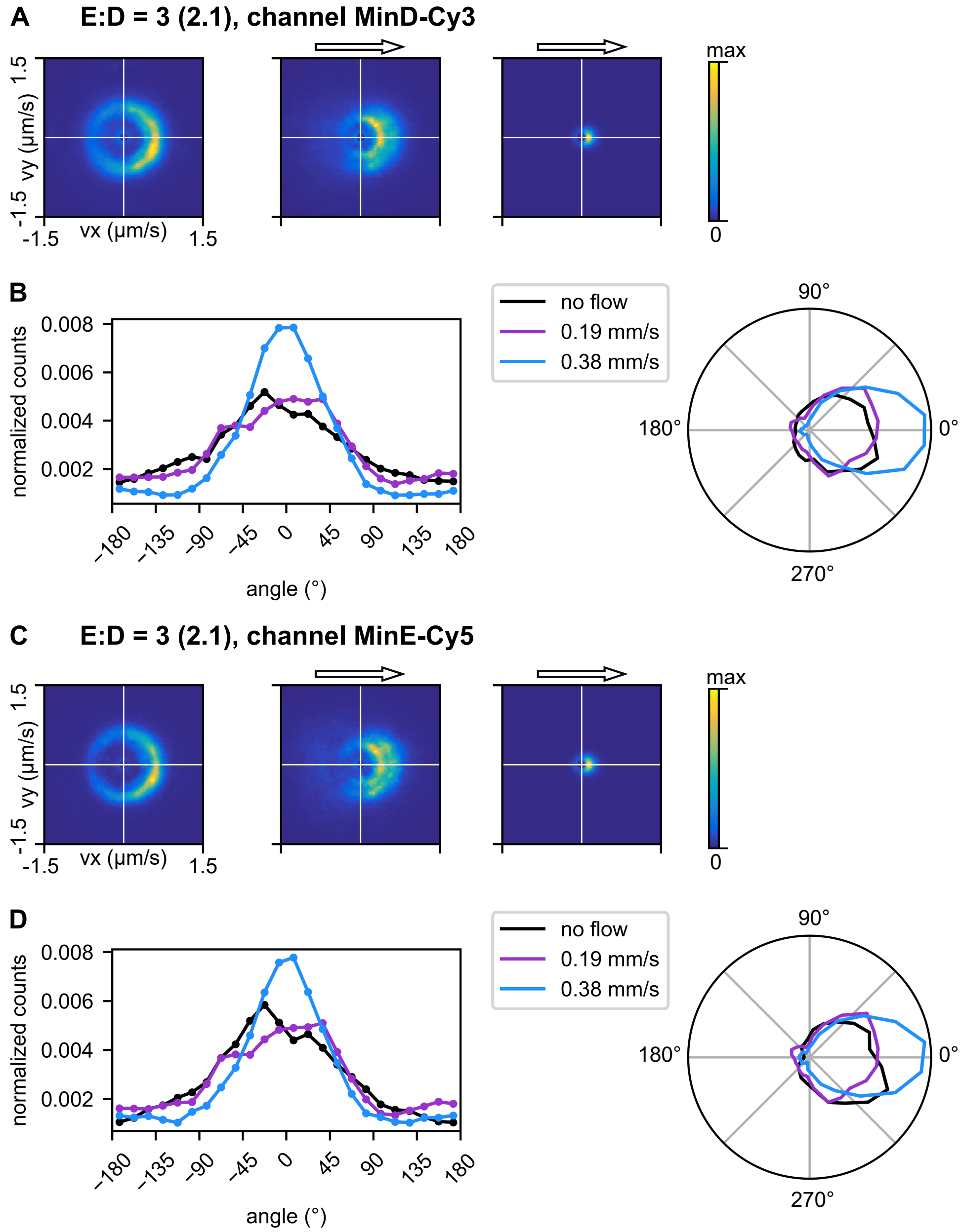


Fig. S17: Analysis for E:D=3 (corrected 2.1), *wt.3.2*, performed with tools presented in Ref.[9] using general smoothing kernel 20 pixels and flow smoothing kernel 50 pixels. Velocity data cropped below 10 % of mean. **A**, **C** Vector counts per segment for no-flow setting and sequentially applied flow rates as given in the box in **B**, represented as 2D histograms (80 bins in $x$-/$y$-direction each), counts per box indicated by color, maximum (yellow) corresponds to maximum bin count for each individual 2D histogram. **B**, **D** 1D histogram and polar histogram representation of counts per direction in bins of 15° for different flow rates.

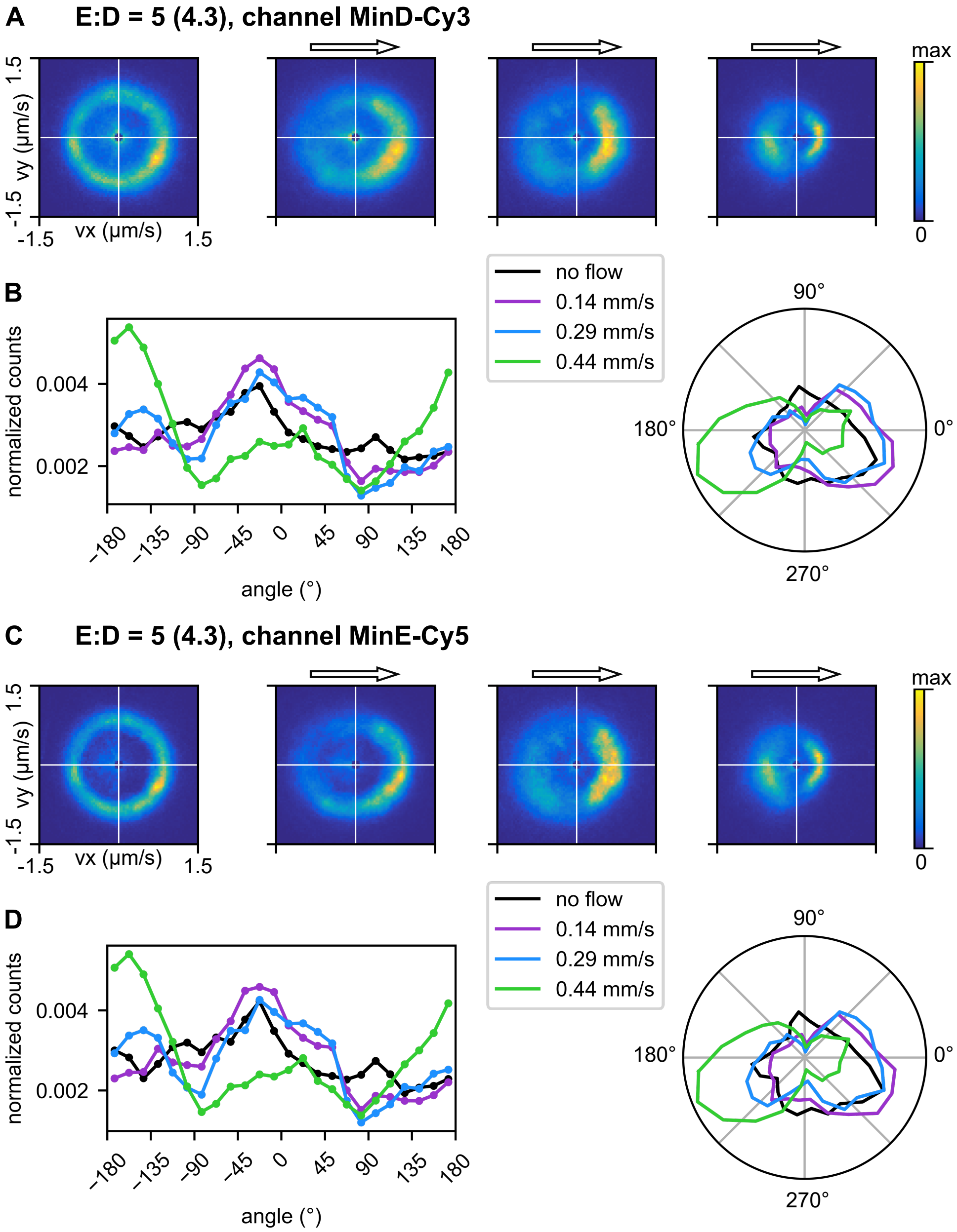


Fig. S18: Analysis for E:D=5 (corrected 4.3), *wt.5.1*, performed with tools presented in Ref.[9] using general smoothing kernel 20 pixels and flow smoothing kernel 50 pixels. Velocity data cropped below 10 % of mean. **A**, **C** Vector counts per segment for no-flow setting and sequentially applied flow rates as given in the box in **B**, represented as 2D histograms (80 bins in $x$-/$y$-direction each), counts per box indicated by color, maximum (yellow) corresponds to maximum bin count for each individual 2D histogram. **B**, **D** 1D histogram and polar histogram representation of counts per direction in bins of 15° for different flow rates.

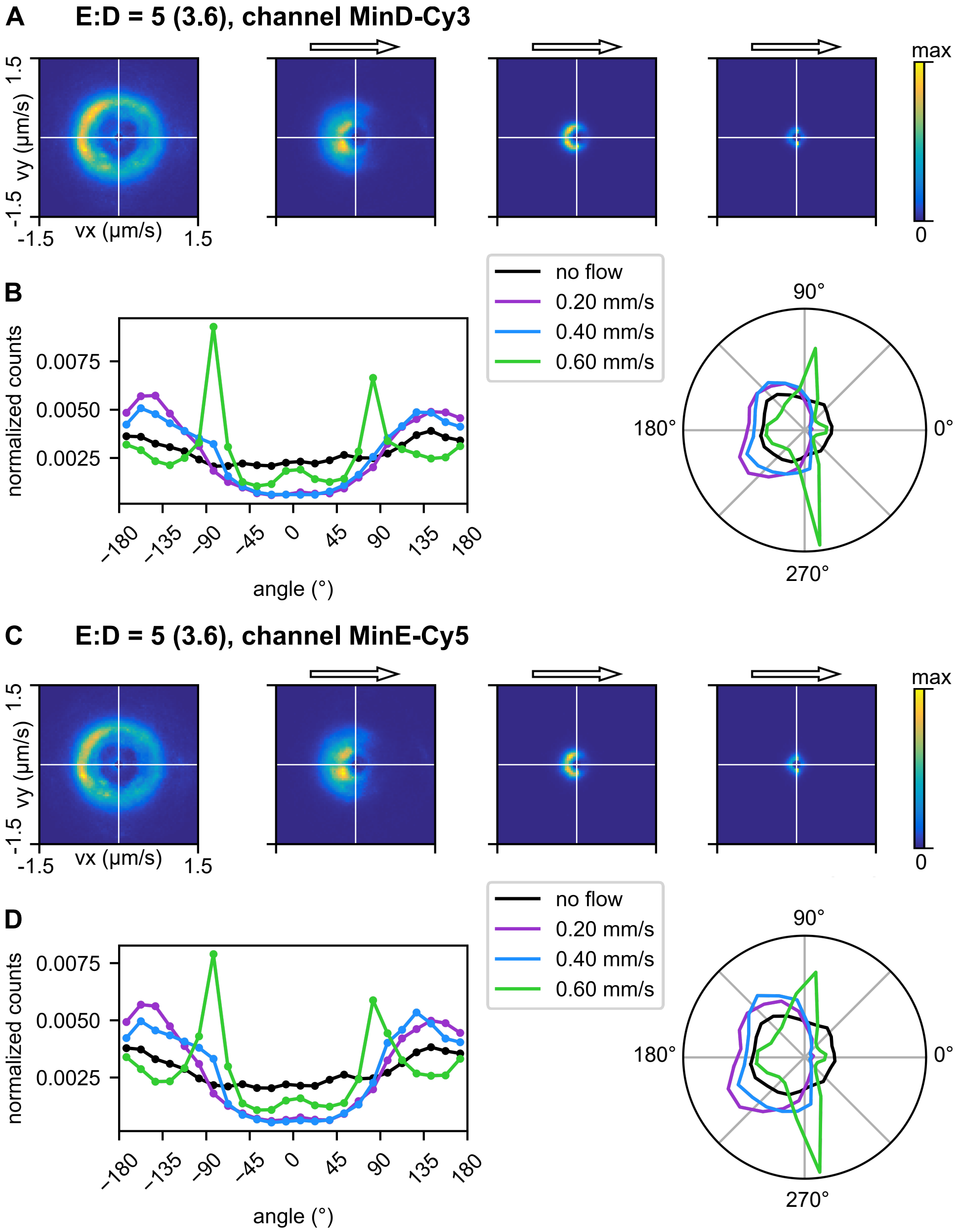


Fig. S19: Analysis for E:D=5 (corrected 3.6), *wt.5.2*, performed with tools presented in Ref.[9] using general smoothing kernel 20 pixels and flow smoothing kernel 50 pixels. Velocity data cropped below 10 % of mean. **A**, **C** Vector counts per segment for no-flow setting and sequentially applied flow rates as given in the box in **B**, represented as 2D histograms (80 bins in $x$-/$y$-direction each), counts per box indicated by color, maximum (yellow) corresponds to maximum bin count for each individual 2D histogram. **B**, **D** 1D histogram and polar histogram representation of counts per direction in bins of 15° for different flow rates.

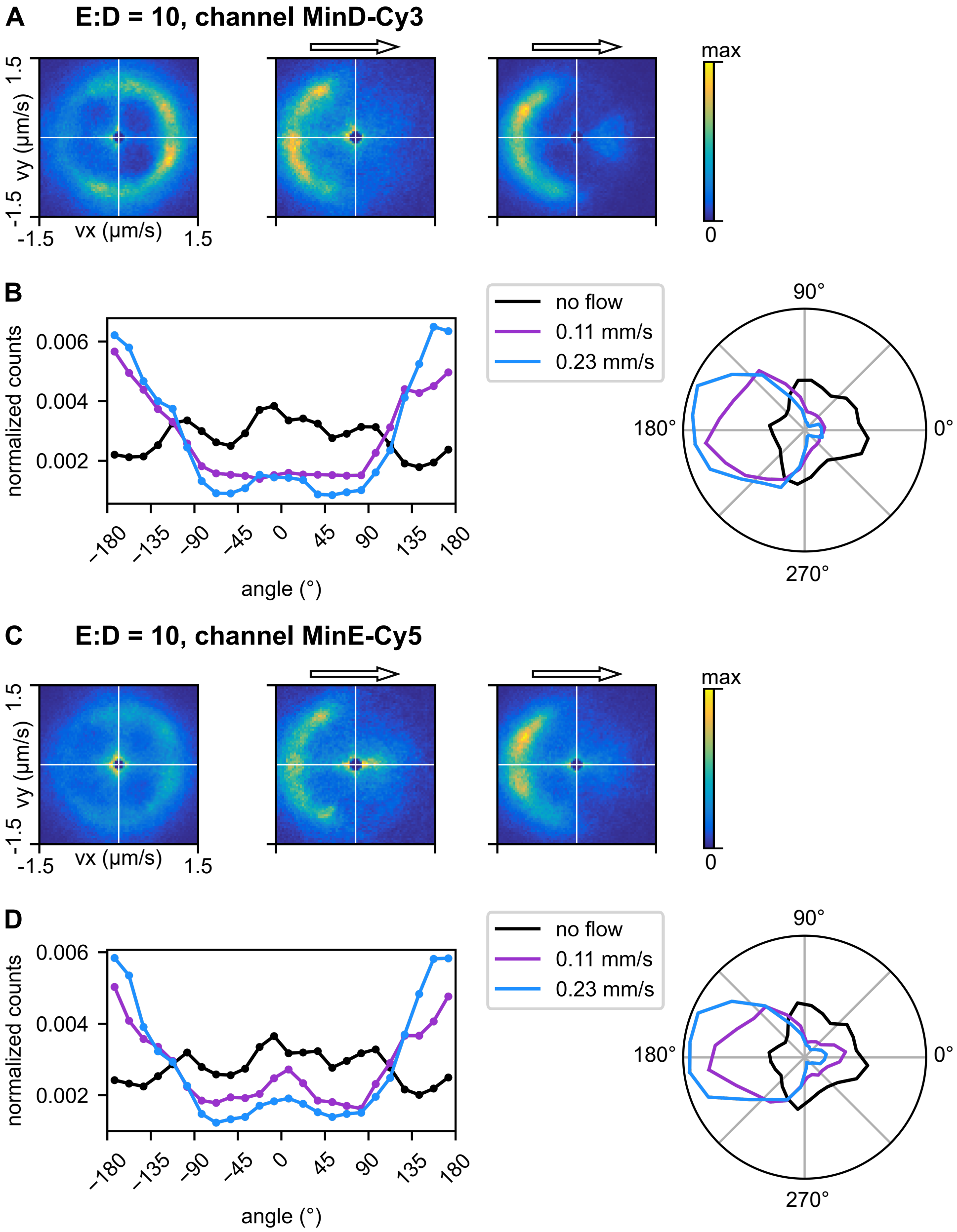


Fig. S20: Analysis for E:D=10, *wt.10.1*, performed with tools presented in Ref.[9] using general smoothing kernel 40 pixels and flow smoothing kernel 50 pixels. Velocity data cropped below 10 % of mean. **A**, **C** Vector counts per segment for no-flow setting and sequentially applied flow rates as given in the box in **B**, represented as 2D histograms (80 bins in $x$-/$y$-direction each), counts per box indicated by color, maximum (yellow) corresponds to maximum bin count for each individual 2D histogram. **B**, **D** 1D histogram and polar histogram representation of counts per direction in bins of 15° for different flow rates.

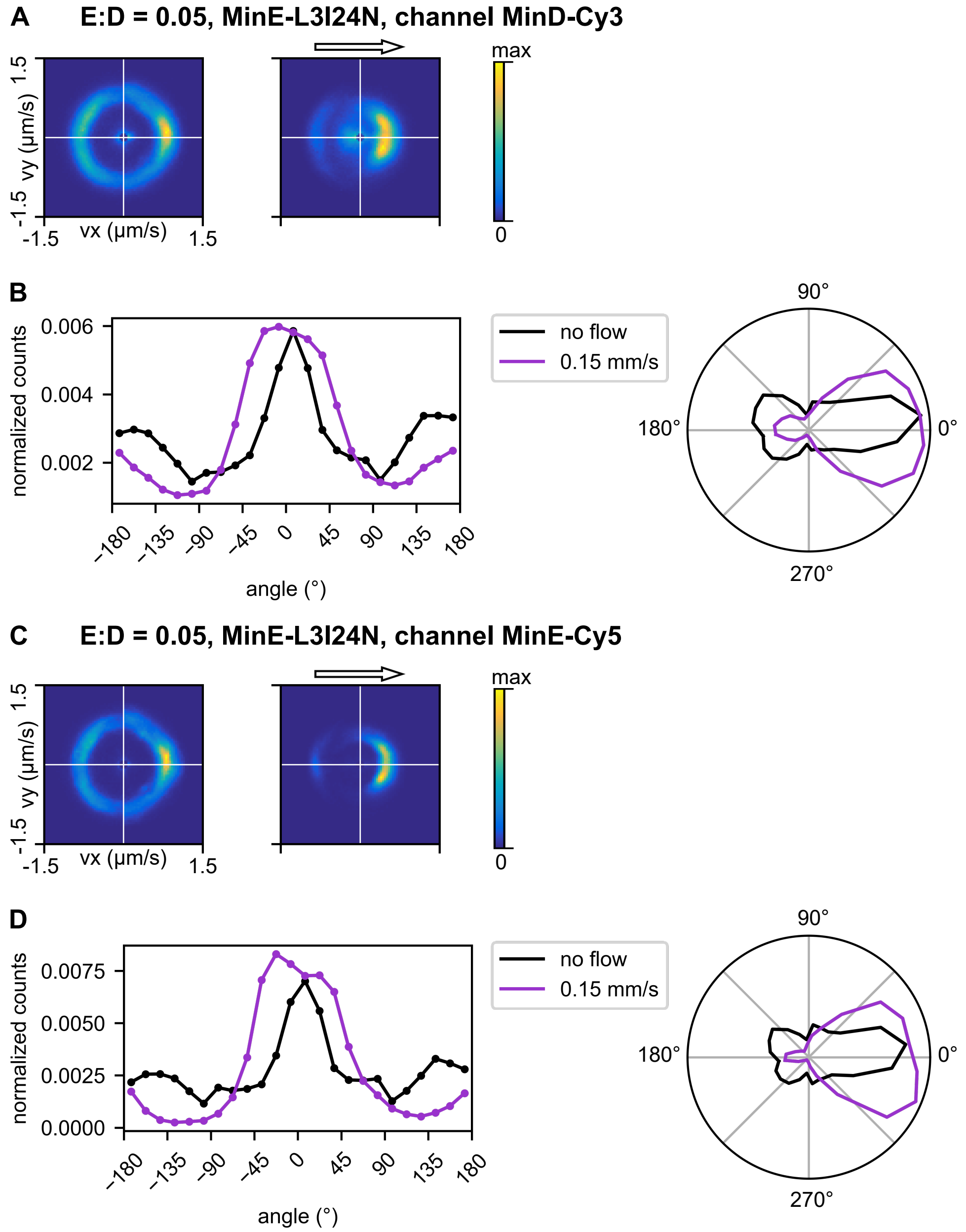


Fig. S21: Analysis for E:D=0.05, MinE-L3I24N, *l3.p05.1*, performed with tools presented in Ref.[9] using general smoothing kernel 15 pixels and flow smoothing kernel 35 pixels. Velocity data cropped below 10 % of mean. **A**, **C** Vector counts per segment for no-flow setting and sequentially applied flow rates as given in the box in **B**, represented as 2D histograms (80 bins in $x$-/$y$-direction each), counts per box indicated by color, maximum (yellow) corresponds to maximum bin count for each individual 2D histogram. **B**, **D** 1D histogram and polar histogram representation of counts per direction in bins of 15° for different flow rates.

#### 5 Supporting movie captions

Six movies are provided along with the main text of the paper. Three of them (movie 1, movie 2, movie 6) contain simulation results, the other three (movie 3, movie 4, movie 5) show exemplary experimental results from the list of acquisitions given in table S3. Movies are provided as mp4 files. The provided movies are:

**Movie 1** (movie_1.mp4) - Simulation result showing downstream propagation, duration 50 s, $n_E$=150 /µm$^2$, $v_\mathrm{f}$=50 µm s$^{-1}$

**Movie 2** (movie_2.mp4) - Simulation result showing upstream propagation, duration 50 s, $n_E$=1000 /µm$^2$, $v_\mathrm{f}$=50 µm s$^{-1}$.

**Movie 3** (movie_3.mp4) - Experimental results for E:D=10, MinE wildtype, without flow, 0.11 mm s$^{-1}$, 0.23 mm s$^{-1}$ and 0.23 mm s$^{-1}$ reversed.

**Movie 4** (movie_4.mp4) - Experimental results for E:D=2 (corrected 1.3), MinE wildtype, without flow, 0.21 mm s$^{-1}$, 0.42 mm s$^{-1}$ and 0.63 mm s$^{-1}$.

**Movie 5** (movie_5.mp4) - Experimental results for E:D=0.05, MinE L3I24N, without flow and 0.15 mm s$^{-1}$.

**Movie 6** (movie_6.mp4) - Simulation result showing upstream to downstream transition, duration 600 s, $n_E$=191.4 /µm$^2$. Flow velocity $v_\mathrm{f}$ decreases linearly from $v_\mathrm{f}$ = 100 µm s$^{-1}$ at $t = 0$ s, to $v_\mathrm{f}$ = 50 µm s$^{-1}$ at $t = 100$ s, remaining at that value until the end of the simulation.

For numerical simulations, unless specified otherwise, parameters given in Table S1 were used, with the units appropriately extended to account for the fact that protein densities have units µm$^{-2}$ in 2D compared to µm$^{-1}$ in a line geometry.

[9] Sabrina Meindlhumer, Jacob Kerssemakers, and Cees Dekker. Quantitative analysis of surface wave patterns of min proteins. *bioRxiv*, 10.1101/2022.04.28.489904, 2022.

[10] Johannes Schindelin, Ignacio Arganda-Carreras, Erwin Frise, Verena Kaynig, Mark Longair, Tobias Pietzsch, Stephan Preibisch, Curtis Rueden, Stephan Saalfeld, Benjamin Schmid, Jean-Yves Tinevez, Daniel James White, Volker Hartenstein, Kevin Eliceiri, Pavel Tomancak, and Albert Cardona. Fiji: an open-source platform for biological-image analysis. 9(7):676–682.

[11] Guido Van Rossum and Fred L. Drake. *Python 3 Reference Manual*. CreateSpace, Scotts Valley, CA, 2009.

[12] Anthony G. Vecchiarelli, Min Li, Michiyo Mizuuchi, and Kiyoshi Mizuuchi. Differential affinities of MinD and MinE to anionic phospholipid influence min patterning dynamics *in vitro*: Flow and lipid composition effects on min patterning. *Molecular Microbiology*, 93(3):453–463, 2014.

[13] Manon C. Wigbers, Fridtjof Brauns, Ching Yee Leung, and Erwin Frey. Flow induced symmetry breaking in a conceptual polarity model. *Cells*, 9(6), 2020.
